# The Scribble/SGEF/Dlg1 complex regulates the stability of apical junctions in epithelial cells

**DOI:** 10.1101/2024.03.26.586884

**Authors:** Agustin Rabino, Sahezeel Awadia, Nabaa Ali, Amber Edson, Rafael Garcia-Mata

## Abstract

SGEF, a RhoG specific GEF, can form a ternary complex with the Scribble polarity complex proteins Scribble and Dlg1, which regulates the formation and maintenance of adherens junctions and barrier function of epithelial cells. Notably, silencing SGEF results in a dramatic downregulation of the expression of both E-cadherin and ZO-1. However, the molecular mechanisms involved in the regulation of this pathway are not known. Here, we describe a novel signaling pathway governed by the Scribble/SGEF/Dlg1 complex. Our results show that an intact ternary complex is required to maintain the stability of the apical junctions, the expression of ZO-1, and TJ permeability. In contrast, only SGEF is necessary to regulate E-cadherin expression. The absence of SGEF destabilizes the E-cadherin/catenin complex at the membrane, triggering a positive feedback loop that exacerbates the phenotype through the repression of E-cadherin transcription in a process that involves the internalization of E-cadherin by endocytosis, β-catenin signaling and the transcriptional repressor Slug.

## Introduction

Most internal organs consist of a monolayer of polarized epithelial cells surrounding a central lumen, which functions to establish a barrier that segregates the internal medium from the outside environment (Rodriguez-Boulan and Macara, 2014). The establishment of cell polarity is regulated by the coordinated action of three highly conserved protein complexes; PAR, Crumbs, and Scribble (Bilder et al., 2003). The Scribble complex, which comprises Scribble, Dlg1 (Discs Large) and Lgl (Lethal Giant Larvae), was originally identified in *Drosophila* as a critical regulator of epithelial polarity (Bilder and Perrimon, 2000; Gateff and Schneiderman, 1974; Mechler et al., 1985; Woods and Bryant, 1991). It was later shown to be involved in the regulation of other cellular processes, including cell-cell adhesion, asymmetric cell division, vesicular trafficking, cell migration, and planar-cell polarity (Elsum et al., 2012). In mammalian cells, the Scribble complex also plays key roles in the regulation of cell adhesion and polarity (Bonello and Peifer, 2019). Importantly, dysregulation of the Scribble complex is commonly observed in human cancers and correlates with tumor progression (Elsum et al., 2012). With no known catalytic activity, the proteins in the Scribble complex are believed to function as scaffolding platforms to recruit other binding partners, including the Rho GTPases and their regulators, such as RhoGEFs and RhoGAPs, to build spatially distinct signaling complexes (Bonello and Peifer, 2019; Iden and Collard, 2008b; Mack and Georgiou, 2014). However, it is not known which downstream signaling pathways are regulated by the Scribble complex.

Previous work from our lab has shown that SGEF (ARHGEF26), a RhoG specific GEF, forms a ternary complex with two members of the Scribble polarity complex: Scribble and Dlg1 (Awadia et al., 2019). SGEF is targeted to the apical junctional complex in a Scribble-dependent fashion and functions in the regulation of barrier function at tight junctions (TJ), and the formation and maintenance of E-cadherin mediated adherens junctions (AJ) (Awadia et al., 2019). Notably, silencing SGEF expression results in a dramatic downregulation of the expression of E-cadherin in epithelial cells. However, the mechanisms by which this novel Scribble/SGEF/Dlg1 complex regulates E-cadherin stability and expression levels are not known.

Here, we describe a novel signaling pathway governed by the Scribble/SGEF/Dlg1 ternary complex. Our results show that the three members of the complex are involved in regulating the stability of both TJ and AJ. At TJ, the ternary complex regulates the expression of ZO-1, and junction permeability. This process relies on the integrity of the ternary complex but is independent of the exchange activity of SGEF. In contrast, only SGEF is involved in regulating E-cadherin expression, and its catalytic activity is essential. Our results demonstrate that silencing SGEF destabilizes the E-cadherin/catenin complex at the membrane which allows for its internalization. This triggers a positive feedback loop, which exacerbates the downregulation of E-cadherin through transcriptional repression, in a process that involves β-catenin signaling and the transcriptional repressor Slug.

## Results

### SGEF, but not Scribble or Dlg1, regulates the expression levels of junctional proteins in epithelial cells

Previously, we have shown that SGEF, a RhoG specific GEF, forms a ternary complex with two members of the Scribble polarity complex, Scribble and Dlg1 (Awadia et al., 2019). We also showed that silencing SGEF expression downregulates the expression of junctional proteins like E-cadherin, afadin, and ZO-1 in epithelial cells (Awadia et al., 2019). However, the specific roles of Scribble and Dlg1 and the relative contribution of each of the ternary complex members to the regulation of E-cadherin and ZO-1 have not been characterized.

To investigate the mechanism and functional consequences for epithelial tissue, we first corroborated our findings by growing MDCK cells on permeable filter inserts. This allows epithelial cells to be exposed to nutrient media from the apical and basolateral sides, which resembles a more physiological state. Similar to what we have shown before, the expression levels of E-cadherin and ZO-1 were drastically reduced in SGEF KD cells when assessed by immunofluorescence (Supp. Figure 1A). This striking phenotype was rescued in SGEF KD cells stably expressing human mNeon-SGEF (Rescue WT; Supp. Figure 1A). Furthermore, we analyzed total cell lysates from confluent monolayers by Western blotting and corroborated that the decrease observed in junctional localization of E-cadherin and ZO-1 in SGEF KD cells was due to reduced protein levels and not re-localization (Supp. Figure 1B, C). In agreement with our previous report, the expression levels of other AJ proteins, such as p120-catenin and β-catenin, were not affected in SGEF KD cells (Figure 1B).

**Figure 1.**
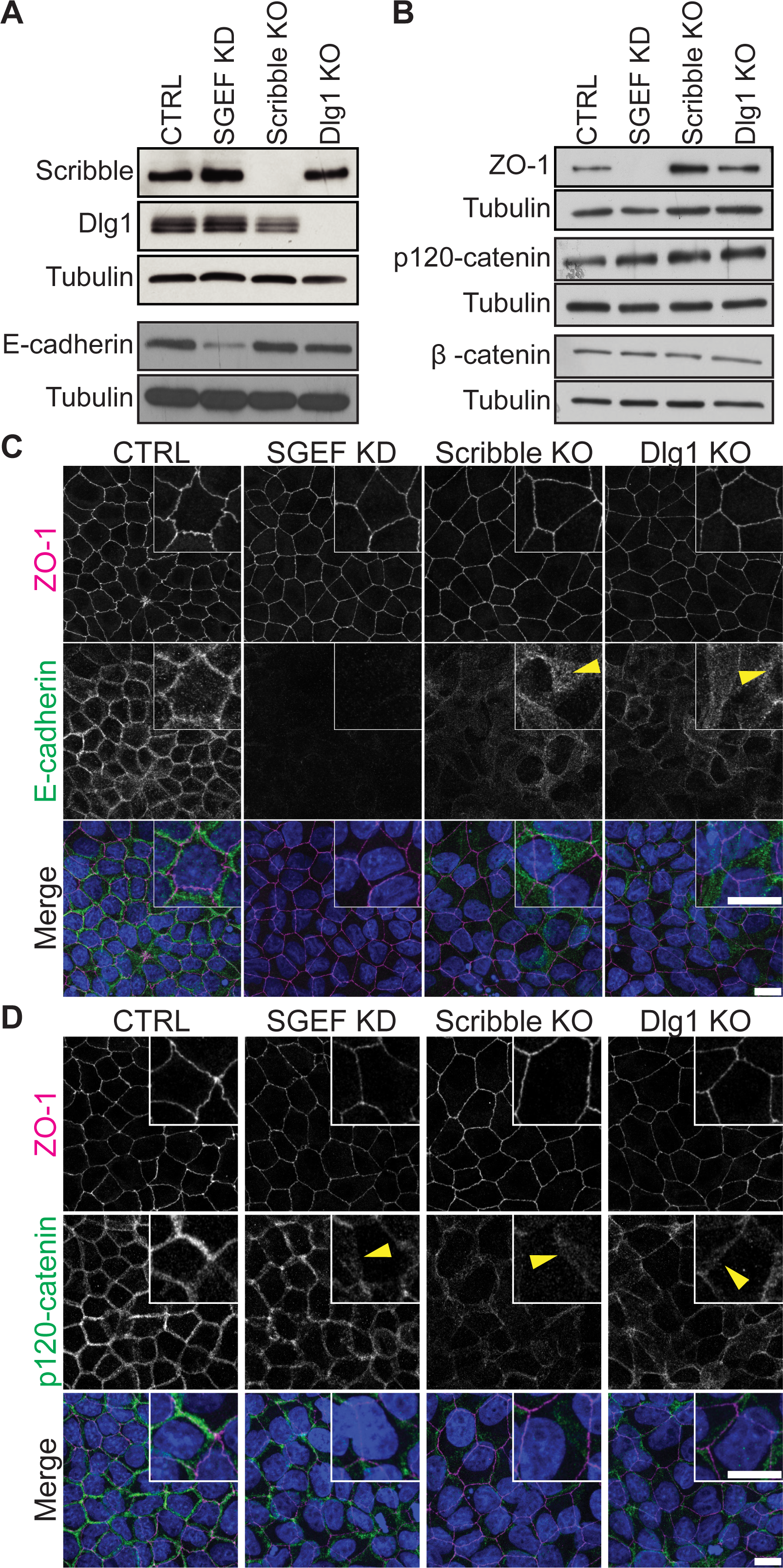
T SGEF, but not Scribble or Dlg1, regulates the expression levels of junctional proteins in epithelial cells. **(A-B)** Total cell lysates from confluent CTRL, SGEF KD, Scribble KO and Dlg1 KO MDCK cells were analyzed by WB for Scribble, Dlg1 and E-cadherin (A); and ZO-1, p120-catenin and β-catenin (B). Tubulin was used as a loading control. **(C-D)** IF showing the distribution of ZO-1 and E-cadherin (C) or ZO-1 and p120-catenin (D) in CTRL, SGEF KD Scribble KO and Dlg1 KO MDCK cells grown in coverslips. The inset shows a 2X zoomed region from the field of view. All images are 3 µm max projections of the subapical domain (ZO-1 signal used for centering). Arrowheads indicate cytosolic localization of E-cadherin. Scale bar 10 µm.

To determine the contribution of Scribble or Dlg1 in the regulation of E-cadherin expression in our model, we generated Scribble and Dlg1 KO MDCK cell lines using CRISPR/Cas9 (Figure 1A). Interestingly, even though Scribble and Dlg1 KO cells showed a slight reduction in E-cadherin expression levels by Western blot, these were not significantly different to CTRL cells (Figure 1A, Supp. Figure 1D). In contrast, the localization of E-cadherin was notably affected in Scribble and Dlg1 KO cells. Both cell lines showed weaker E-cadherin staining in general, which was less defined at the cell-cell junctions and had increased localization at punctate structures in the interior of the cells (Figure 1C, arrowheads). Other AJ proteins like p120-catenin and β-catenin did not show changes at the total protein level (Figure 1B) but showed a similar re-localization as E-cadherin (Figure 1D, Supp. Figure 1E, arrowheads). These results suggest that, of the three complex members, only SGEF plays a key role in the regulation of E-cadherin expression.

Since Scribble and Dlg1 have also been associated to TJ formation, we analyzed the localization of the TJ marker ZO-1 in Scribble KO and Dlg1 KO cells (Elsum et al., 2013; Ivanov et al., 2010; Yates et al., 2013). Interestingly, both Scribble and Dlg1 KO cells phenocopied SGEF KD cells at the TJ level, showing a decreased amount of ZO-1 at the junctions, and a reorganization of TJs into a more linear phenotype than in CTRL MDCK cells, which typically display curvilinear phenotype (Figure 1C, Supp. Figure 2A-B). Interestingly, the decrease in ZO-1 intensity at the junctions correlated with a decrease in proteins levels only in SGEF KD cells, suggesting that in Scribble and Dlg1 KO cells ZO-1 may be redistributed off the junctions (Figure 1B, Supp. Fig. 2C). To characterize this phenotype in depth, we analyzed the shape of cells defined by the TJs contour (area, circularity, axial and feret ratio), as well as the tortuosity index. To better represent all the data collected we used a principal component analysis (PCA) (Supp. Figure 2D) in which the main components (PC1 and PC2) describe the relationship with the other parameters through a correlation table (Supp. Figure 2E). The results from the PCA analysis showed that both Scribble KO and Dlg1 KO cells phenocopy SGEF KD cells in terms of junctions’ shape and TJ tortuosity index. The three cell lines (Scribble KO, Dlg1 KO and SGEF KD) had a bigger apical area, were more regular in shape, as seen by the circularity, axial and feret ratios all being close to a value of 1, and had TJs that were more linear/straight (less tortuous).

To assess the functional implications of these phenotypes, we analyzed the ability of CTRL, SGEF KD, Scribble KO and Dlg1 KO cells to form a monolayer and establish an impermeable barrier. We measured the Trans Epithelial Electrical Resistance (T.E.E.R.) which measures the charge-selective permeability of small solutes in confluent monolayers grown on semi-permeable filters. T.E.E.R also provides an indication of TJ barrier function (Anderson and Van Itallie, 2009). We have reported before that SGEF KD cells have a significant reduction in the T.E.E.R. compared to CTRL MDCKs (Awadia et al., 2019). Here, our results also showed a decrease in T.E.E.R. in both Scribble KO and Dlg1 KO. T.E.E.R. was almost 50% lower in both Scribble and Dlg1 KO cells compared to CTRL cells, similar to the reduction observed for SGEF KD cells (Supp. Figure. 2F).

To summarize, the three members of the Scribble/SGEF/Dlg1 complex play a similar role in the regulation of tight junctions’ architecture and barrier function, but only SGEF plays a significant role in the regulation of E-cadherin and ZO-1 protein expression.

### SGEF’s guanine exchange activity is required to rescue E-cadherin levels in SGEF KD cells

Our previous results demonstrated that a catalytic dead (CD) mutant of SGEF was not able to rescue the downregulation of E-cadherin (Awadia et al., 2019). This suggested that SGEF’s guanine exchange activity was necessary to rescue E-cadherin levels in SGEF KD cells (Awadia et al., 2019). Here, we designed two new constructs to better understand the role of SGEF targeting by the ternary complex and its catalytic activity in the regulation of E-cadherin levels. The first construct encoded the catalytic domain of SGEF fused to GFP (DH-PH-GFP). This construct lacks any known targeting information and cannot bind to Scribble or Dlg1. The second construct encoded the catalytic domain of SGEF fused to the VSV-G protein (VSV-G-DH-PH-GFP) which targets it to the basolateral membrane independently of Scribble and Dlg1 (Figure 2A) (Keller et al., 2001). Using these constructs in SGEF KD cells we aimed to address two potential scenarios: a) SGEF activity alone is sufficient to rescue E-cadherin expression, regardless of its localization; b) SGEF activity localized at the basolateral membrane is needed to rescue E-cadherin expression. To distinguish between these two possibilities, we stably expressed each construct in SGEF KD MDCK cells and analyzed their ability to rescue E-cadherin levels by Western blot and immunofluorescence. As controls, we also analyzed the previously characterized Rescue WT and Rescue CD stable cell lines (Awadia et al., 2019). Our results showed that E-cadherin levels were significantly rescued only when the catalytic activity was targeted to the basolateral membrane (Rescue WT, and VSV-G-DH-PH) (Figure 2B, C). Immunofluorescence staining showed that targeting the catalytic activity of SGEF to the membrane using VSV-G not only restored the expression levels of E-cadherin, but also its proper junctional localization (Figure 2D, E). In contrast, cells expressing DH-PH-GFP showed a slight but not significant increase in E-cadherin protein levels, and a small increase in junctional localization of E-cadherin (Figure 2B-E). We believe this could be a result of a non-specific global increase in RhoG activation. These results support our hypothesis that at least one of the functions of the complex is to target SGEF activity to the correct cellular localization, where its catalytic activity is required for regulation of E-cadherin turnover.

**Figure 2.**
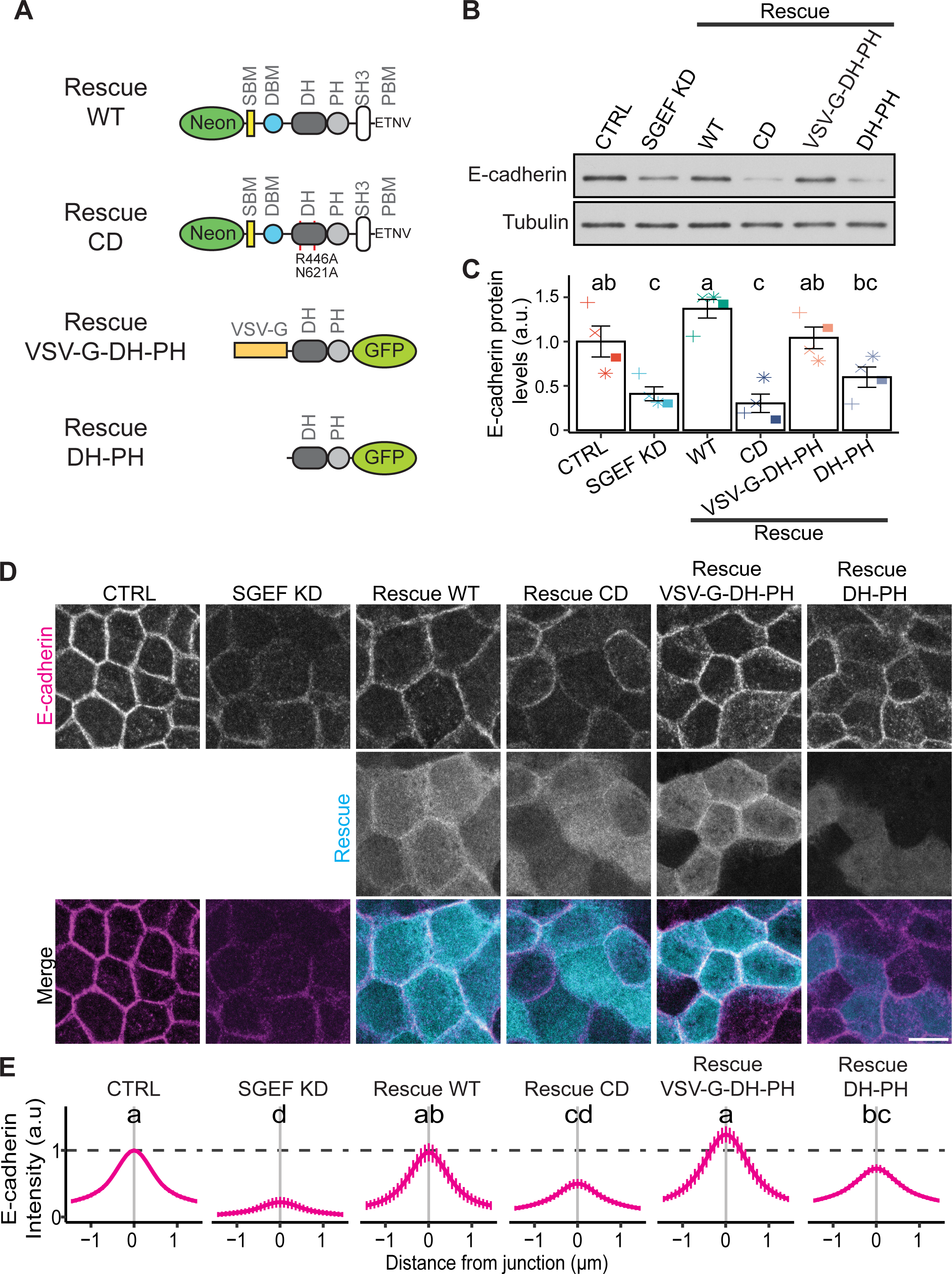
SGEF catalytic activity is required at the basolateral membrane for E-cadherin junctional stability. **(A)** Schematic representation of SGEF constructs used in this study. **(B)** Total cell lysates from CTRL, SGEF KD, and SGEF KD cells rescued with mNeon-SGEF WT, mNeon-SGEF CD, VSV-G-DH-PH-GFP, or DH-PH-GFP were probed for E-cadherin. Tubulin was used as a loading control. **(C)** Quantification and ANOVA analysis from WB results, letters indicate Tukey post-hoc significant differences between groups (n=4). **(D)** IF of CTRL, SGEF KD, and SGEF KD cells rescued with mNeon-SGEF WT, mNeon-SGEF CD, VSV-G-DH-PH-GFP, or DH-PH-GFP, grown in cover slips and stained for E-cadherin. All images are 3 µm max projections of the subapical domain (Cortical Actin signal used for centering). **(E)** Quantification of E-cadherin intensity measured from a perpendicular line across the junctions. Letters indicate significant differences in the intensity of E-cadherin between cell lines (Tukey post-hoc analysis) (n=5; >200 junctions/condition across all experiment). Scale bar 10µm.

### E-cadherin downregulation is not dependent on protein degradation

We then investigated the molecular mechanisms controlling the SGEF-dependent downregulation of E-cadherin. Our initial hypothesis was that E-cadherin degradation was enhanced when SGEF expression was silenced, and inhibiting the proteolytic system involved would rescue E-cadherin levels. To test our hypothesis, we used the commercially available inhibitors MG132 and chloroquine to inhibit proteasomal and lysosomal degradation respectively and analyzed the expression levels of E-cadherin by Western blotting and immunofluorescence. Our results showed a small but reproducible increase in E-cadherin levels in cells treated with the proteasome inhibitor MG132 (Figure 3A-B). In contrast, inhibition of lysosomal degradation showed no appreciable difference. However, even after overnight treatment with MG132, E-cadherin levels were still almost 50% lower in SGEF KD cells when compared to CTRL, suggesting that inhibiting the proteasome is not sufficient to restore normal levels of E-cadherin in the absence of SGEF (Figure 3B). This was supported by a post-hoc Tukey analysis, which showed that overall, SGEF KD cells treated with the two inhibitors are statistically the same as non-treated cells (Figure 3B). Additionally, immunofluorescence analysis showed no major changes in the localization of E-cadherin when the proteasome and lysosome systems were inhibited (Figure 3C, D). Overall, our results suggest that the downregulation of E-cadherin observed in SGEF KD cells cannot be attributed to an increase in protein degradation.

**Figure 3.**
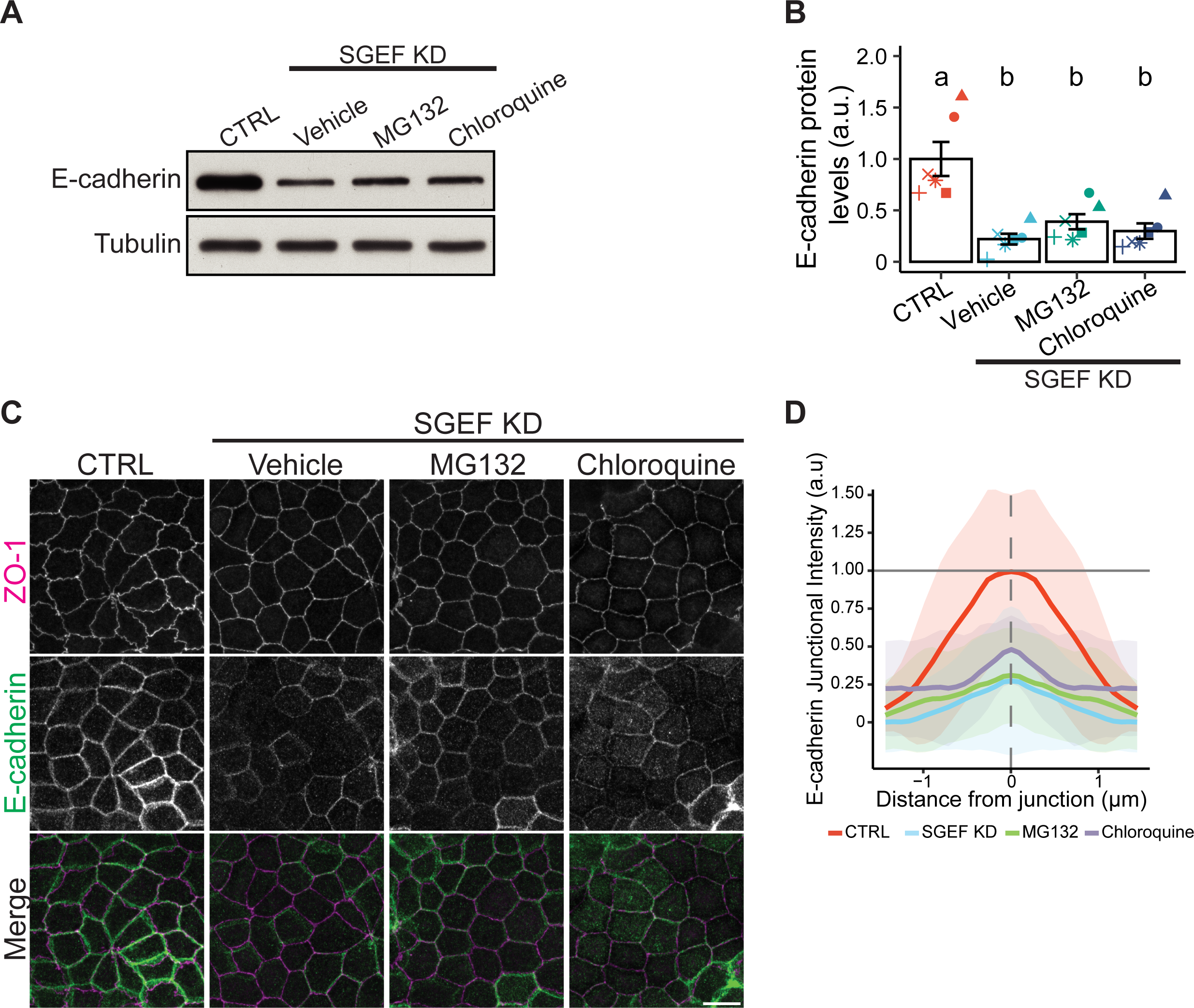
E-cadherin downregulation in SGEF KD cells is not mediated by protein degradation. **(A)** Total cell lysates from confluent CTRL (non-treated) and SGEF KD MDCK cells treated with the indicated inhibitors for 16 h were analyzed for E-cadherin expression by WB. Tubulin was used as a loading control. **(B)** Quantification and ANOVA analysis from WB results, letters indicate Tukey post-hoc significant differences between groups (n=6). **(C)** IF of endogenous ZO-1 and E-cadherin from CTRL (non-treated) and SGEF KD cells after incubation with the indicated inhibitors. **(D)** Quantification of E-cadherin intensity measured from a perpendicular line across the junctions (n=1; >30 junctions/condition, shaded area represents SD). All images are 3 µm max projections of the subapical domain (ZO-1 signal used for centering). Scale bar 10 µm.

### SGEF guanine exchange activity regulates E-cadherin transcription in a Slug mediated fashion

Since the E-cadherin downregulation in SGEF KD cells could not be explained by an increase in protein degradation, we analyzed E-cadherin mRNA levels in CTRL, SGEF KD, Rescue WT, and Rescue CD cells using real time PCR (Figure 4A). Our results showed that E-cadherin mRNA levels were significantly downregulated in SGEF KD cells, and the re-expression of SGEF WT (Rescue WT) restored E-cadherin mRNA levels back to normal. In contrast, re-expression of SGEF CD (Rescue CD) was not able to rescue E-cadherin mRNA levels, underscoring the essential role of SGEF’s catalytic activity in the transcriptional regulation of E-cadherin.

**Figure 4.**
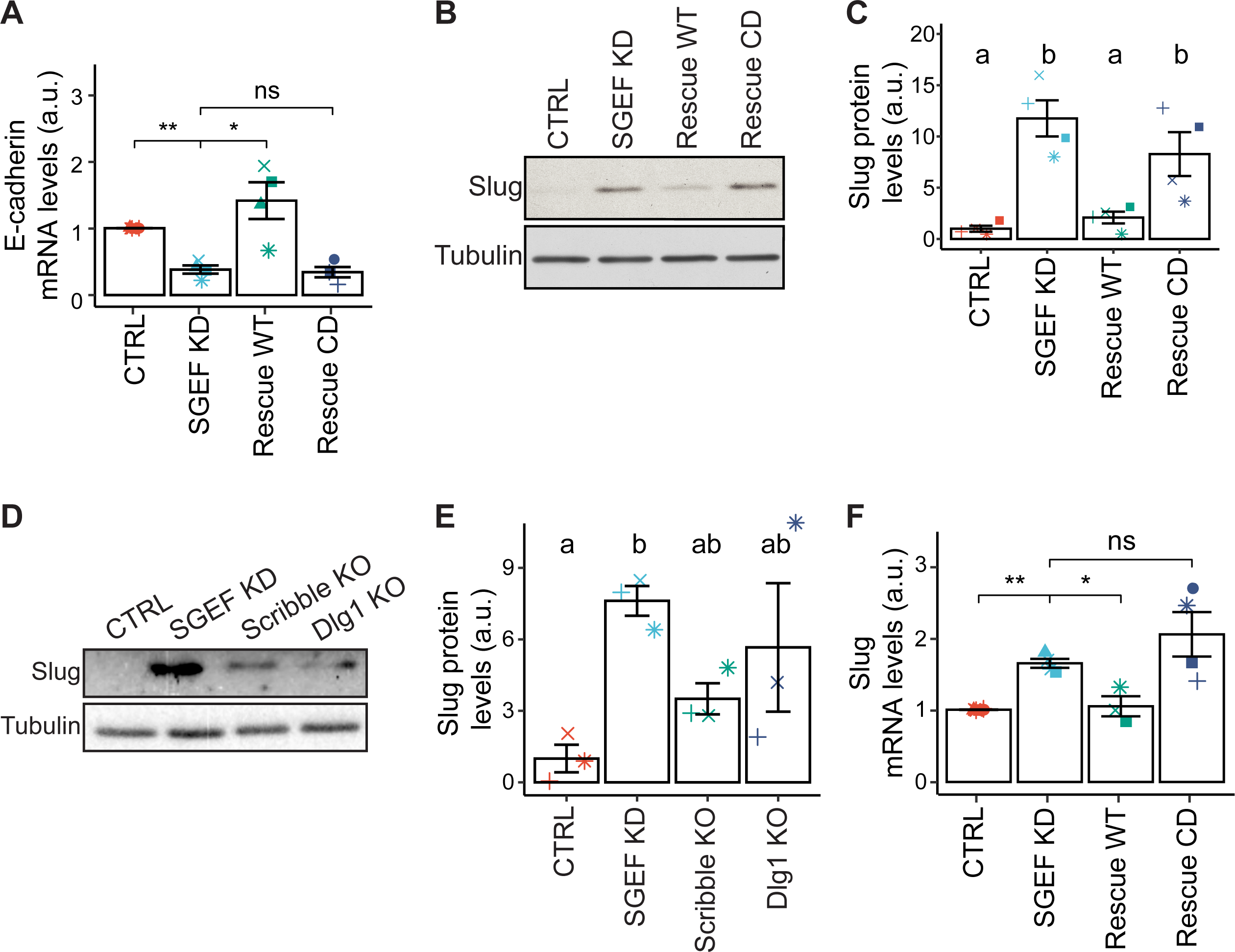
The guanine exchange activity of SGEF regulates E-cadherin transcription in a Slug mediated fashion. **(A)** qPCR analysis of E-cadherin transcript levels in confluent CTRL, SGEF KD, Rescue WT and Rescue CD MDCK cells. **(B)** Slug protein levels were analyzed by WB in confluent CTRL, SGEF KD, Rescue WT and Rescue CD MDCK cells. Tubulin was used as a loading control. **(C)** Quantification and ANOVA analysis from WB data in (B). Letters indicate Tukey post-hoc significant differences between groups (n=4). **(D)** Slug protein levels were analyzed by WB in confluent CTRL, SGEF KD, Scribble KO and Dlg1 KO MDCK cells. Tubulin was used as a loading control. **(E)** Quantification and ANOVA analysis from WB data in (D). Letters indicate Tukey post-hoc significant differences between groups (n=3). **(F)** qPCR analysis of Slug transcript levels in CTRL, SGEF KD, Rescue WT and Rescue CD cells.

E-cadherin transcriptional downregulation has been widely studied in the context of epithelial to mesenchymal transition (EMT), and it is known that the Snail and Zeb families of transcriptional repressors play a key role in this process (Batlle et al., 2000; Cano et al., 2000; Comijn et al., 2001; Conacci-Sorrell et al., 2003). To understand the nature of E-cadherin transcriptional downregulation in SGEF KD cells, we analyzed the expression levels of the transcription factors known to be associated with E-cadherin repression. As a positive control, we treated the cells with hepatocyte growth factor (HGF), which is known to induce the expression of the Snail family members and induce EMT in MDCK cells (Lee et al., 2011; Leroy and Mostov, 2007). We found that Slug, a member of the Snail family of transcriptional repressors, was drastically upregulated (up to 10-fold) in SGEF KD cells (Figure 4B-C, Supp. Figure 3A). Other known E-cadherin repressors, including Snail and ZEB1 showed no change compared to CTRL (Supp. Figure 3B, C). In addition, we immunoblotted for other EMT markers, like the intermediate filament protein vimentin and N-cadherin, and saw no change in their expression between CTRL and SGEF KD cells (Supp. Figure 3D, E). These results suggest that SGEF KD cells have not undergone complete EMT.

We then used our previously established cell lines to understand the contributions of the SGEF’s catalytic activity and the other complex members in the regulation of Slug. Similar to what we observed for E-cadherin, the catalytic activity of SGEF was essential for the regulation of Slug protein levels, as they were restored to CTRL levels only when SGEF WT was re-expressed (Rescue WT), but not with catalytically inactive SGEF (Rescue CD) (Figure 4B). ANOVA and Tukey post-hoc analysis showed a clear significant difference between these groups (Figure 4C). Interestingly, we saw a small but reproducible increase in Slug levels when we knocked out Scribble and Dlg1 (Figure 4D). Even though these differences were not significantly different from CTRL (Figure 4E), the increase in Slug correlates with the small decrease in E-cadherin observed in Scribble and Dlg1 KO cells (Sup. Figure 1D) and may reflect an intermediate less severe phenotype.

Slug can be regulated both at the mRNA and protein level (Kim et al., 2014; Moon et al., 2021; Vallin et al., 2001; Wang et al., 2009). Here, we sought to answer what was the nature of Slug upregulation in SGEF KD cells. Our results showed that the changes observed in Slug protein levels closely correlate with changes in Slug mRNA, with a significant increase of Slug mRNA in SGEF KD cells, suggesting that Slug expression is regulated at the level of transcription (Figure 4F). Slug mRNA levels are rescued by re-expressing SGEF WT but not a catalytic dead mutant, confirming the role of SGEF catalytic activity in regulating Slug expression through transcription.

In summary, we showed that E-cadherin is downregulated at the transcriptional level in SGEF KD cells, and that this decrease in E-cadherin mRNA levels is accompanied by the upregulation of the transcriptional repressor Slug. The regulation of both E-cadherin and Slug levels depends on the catalytic activity of SGEF.

### Silencing of Slug in SGEF KD cells rescues E-cadherin protein levels

To confirm that the downregulation of E-cadherin in SGEF KD cells is mediated by the upregulation of Slug levels, we stably silenced the expression of Slug in SGEF KD cells using lentivirus. We predicted that silencing Slug in SGEF KD cells should rescue E-cadherin expression. Our results showed that Slug expression was efficiently silenced in the stable SGEF/Slug dKD (double KD) cells, with Slug expression levels comparable to CTRL MDCK cells (Figure 5A, B). Importantly, when compared to SGEF KD cells, E-cadherin levels in the double KD cells were rescued to a level that was even higher than those of CTRL cells (Figure 5A, C). We also analyzed the distribution of endogenous E-cadherin in CTRL, SGEF KD, and SGEF/Slug dKD cells by immunofluorescence. As seen in Figure 5E, SGEF/Slug dKD cells showed rescue of E-cadherin. Interestingly, we noticed that E-cadherin staining does not localize exclusively to the lateral junctions in the double KD and was found also in the cytosol and the basal membrane of the cell (Figure 5E, arrowheads).

**Figure 5.**
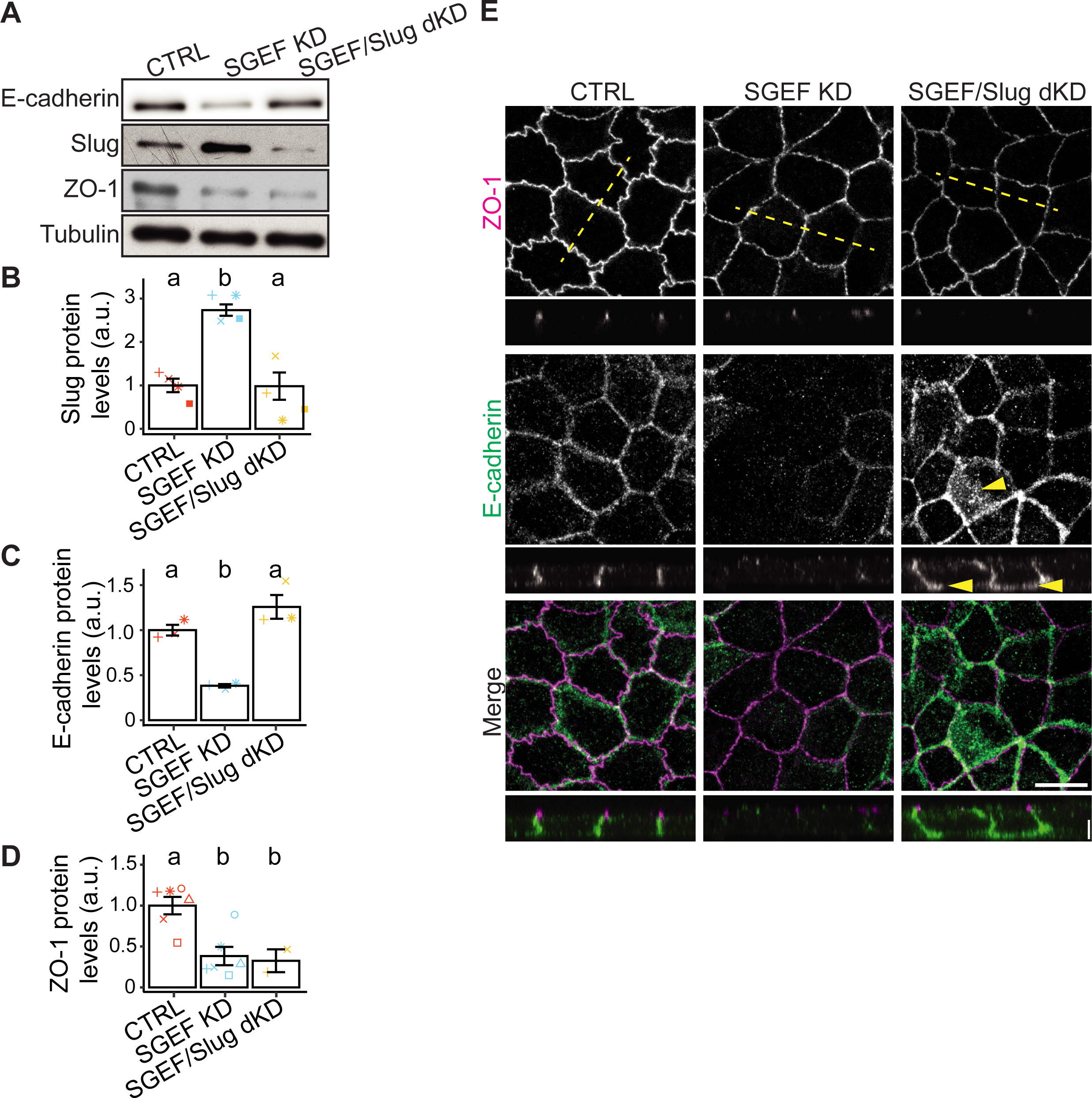
Silencing Slug in SGEF KD cells rescues E-cadherin protein levels. **(A)** Total cell lysates from confluent CTRL, SGEF KD, and SGEF/Slug dKD MDCK cells were analyzed by WB for E-cadherin, Slug and ZO-1. Tubulin was used as a loading control. **(B-D)** Quantification and ANOVA analysis from WB data in (A). Letters indicate Tukey post-hoc significant differences between groups. **(E)** IF of endogenous ZO-1 and E-cadherin in confluent CTRL, SGEF KD, and SGEF/Slug KD MDCK cells. The yellow lines indicate the section used for XZ cross-section shown in the bottom panels. All images are 3 µm max projections of the subapical domain (ZO-1 signal used for centering). Scale bar 10µm (XY) and 5µm (XZ).

Surprisingly, Slug KD did not rescue the expression of ZO-1, which is also downregulated in SGEF KD cells, as we have previously shown (Awadia et al., 2019) (Figure 5A, D). Supporting the results of the Western blot, immunofluorescence staining shows that there is no significant recovery in the ZO-1 signal at junctions in the double KD cells (Figure 5E). Analysis of ZO-1 mRNA levels by qPCR shows a significant decrease in SGEF KD cells when compared to CTRL cells (Supp. Figure 4A). These results suggest that Slug is not involved in the downregulation of ZO-1 observed in SGEF KD cells.

To better characterize the distribution of E-cadherin and ZO-1 along the basolateral membrane, we quantified their intensities along the confocal Z-stack. To register the measurements from different experiments, we normalized the values and aligned all junctions to the brightest Z-plane of ZO-1. Our results showed a clearly defined peak for ZO-1 in CTRL cells, which was located approximately 1 µm apically of the peak for E-cadherin (Supp. Figure 4B, red lines). SGEF KD cells showed a marked decrease in both intensity profiles, supporting our Western Blot results (Supp. Figure 4B, cyan lines). In contrast, SGEF/Slug dKD cells showed similar E-cadherin intensity profile than that in CTRL cells, confirming an overall rescue. However, the distribution of E-cadherin displayed increased variability along the basolateral membrane (Supp. Figure 4B, orange lines, see SEM marked by dashed lines). This correlates with the redistribution of E-cadherin signal from the lateral membrane in CTRL cells to the lateral and basal membrane in the double KD cells (Figure 5E, arrowheads). Interestingly there was a slight rescue of ZO-1 when we quantified the immunofluorescence images (Supp. Figure 4B). However, the overall ZO-1 protein levels remained significantly different to CTRL cells as seen in Figure 5A and D.

Overall, our results demonstrate that E-cadherin downregulation in SGEF KD cells is driven by an upregulation of Slug and suggests that ZO-1 levels are regulated through a different mechanism. In addition, they suggest that rescuing E-cadherin levels without SGEF is not sufficient to restore its proper localization.

### β-catenin signaling regulates E-cadherin and ZO-1 downregulation in SGEF KD cells by two independent pathways

The β-catenin signaling pathway has been reported to be a positive stimulator of EMT, increasing cell invasion and metastasis (Novak and Dedhar, 1999). Interestingly, β-catenin activation functions as a transcriptional co-activator for the T-cell factor (TCF)/Lymphoid enhancer-binding factor (LEF) transcription factors (Valenta et al., 2012). To determine if the β-catenin pathway was responsible for the upregulation of Slug levels observed in SGEF KD cells, we treated cells with iCRT3, an inhibitor of the β-catenin responsive transcription (CRT) (Gonsalves et al., 2011). Importantly, iCRT3 inhibits β-catenin transcriptional coactivator activity but has no effect on its interaction with E-cadherin (Gonsalves et al., 2011). Our results show that treating SGEF KD cells with iCRT3 reduced Slug expression to CTRL levels (Figure 6A, B). In addition, E-cadherin expression was rescued to levels that were higher than those in CTRL cells (Figure 6A, C). Surprisingly, and in contrast to the results obtained with the SGEF/Slug dKD cells, the decrease in ZO-1 levels observed in SGEF KD cells was also restored to CTRL levels after iCRT3 treatment (Figure 6A, D). This suggests that β-catenin signaling pathway is regulating ZO-1 expression via a transcriptional repressor different than Slug.

**Figure 6.**
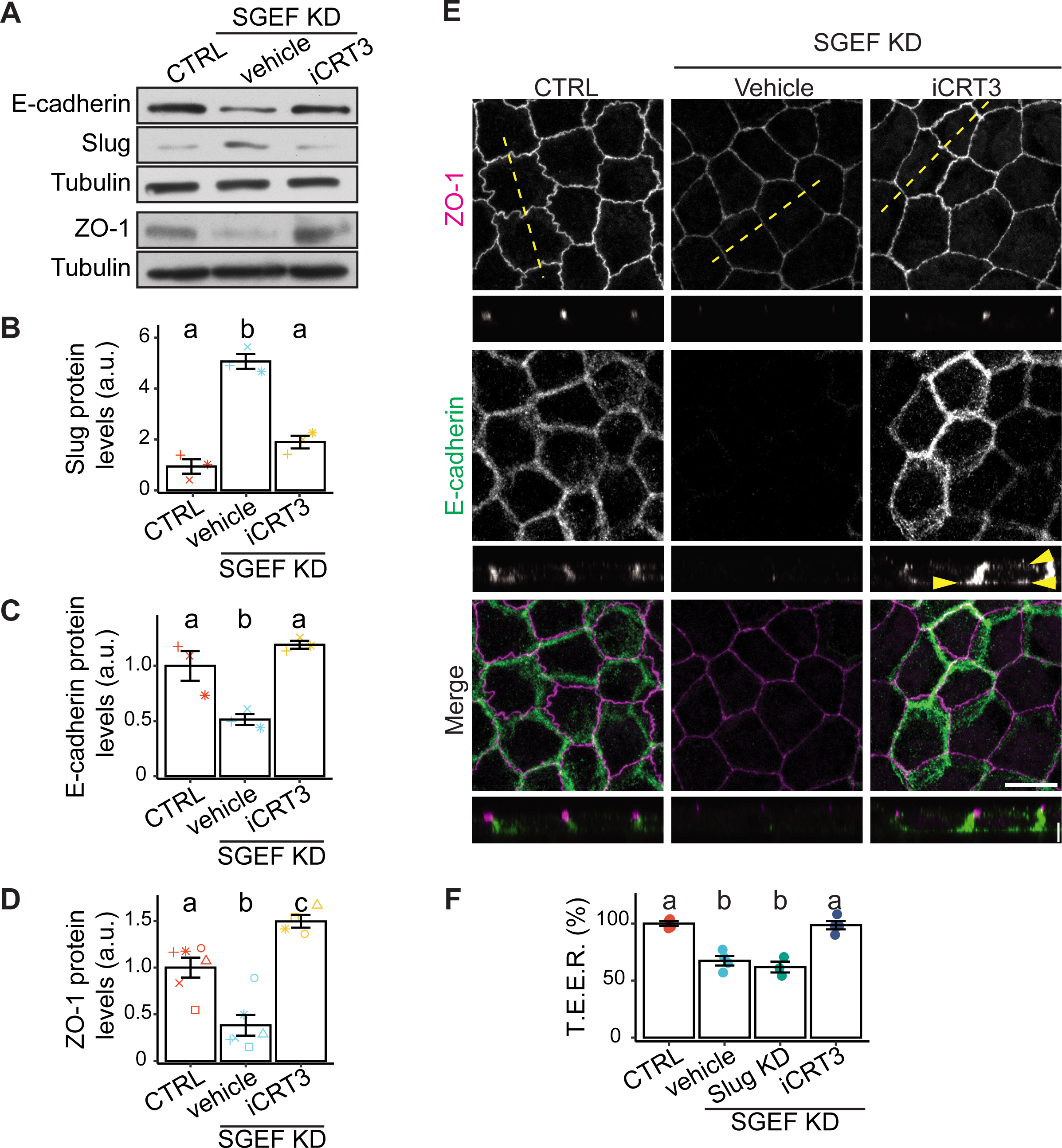
Inhibition of β-catenin signaling pathway rescues E-cadherin and ZO-1 downregulation in SGEF KD cells. **(A)** Total cell lysates from confluent CTRL, SGEF KD, and iCRT3 treated SGEF KD MDCK cells were analyzed by WB for E-cadherin, Slug and ZO-1. Tubulin was used as a loading control. **(B-D)** Quantification and ANOVA analysis from WB data in (A). Letters indicate Tukey post-hoc significant differences between groups. **(E)** IF of endogenous ZO-1 and E-cadherin confluent CTRL (non-treated), SGEF KD, and iCRT3 treated SGEF KD MDCK cells. The yellow lines indicate the region displayed in the XZ cross-section. All images are 3 µm max projections of the subapical domain (ZO-1 signal used for centering) **(F)** T.E.E.R. was measured in CTRL, SGEF KD, SGEF/Slug dKD, and iCRT3 treated SGEF KD MDCK cells grown in permeable filters. The graph represents the difference in electrical resistance as a percentage of CTRL cells (n=4). Scale bar 10µm (XY) and 5µm (XZ).

We then compared the localization of endogenous E-cadherin and ZO-1 between CTRL, untreated SGEF KD, and iCRT3 treated SGEF KD cells. Our results showed that E-cadherin intensity is rescued when SGEF KD cells are treated with iCRT3. In iCRT3-treated SGEF KD cells E-cadherin localized both at the lateral and basal membranes, as opposed to the typical lateral membrane localization observed in CTRL cells (Figure 6E, arrowheads). The redistribution of E-cadherin was similar to that observed in SGEF/Slug dKD cells (Figure 5E). In addition, SGEF KD cells treated with iCRT3 showed increased ZO-1 intensity and more cytosolic E-cadherin (Figure 6E, XZ cross section).

We measured the relative distribution and intensity of the junctional proteins ZO-1 and E-cadherin along the Z-axis (Supp. Figure 4C), as we did in Supp. Figure 4B. Similar to SGEF/Slug dKD cells, the E-cadherin intensity profile in SGEF KD cells treated with iCRT3 (orange lines) was comparable to CTRL cells. However, the quantification displayed more variability along the basolateral membrane (Supp. Figure 4C, see SEM marked by dashed lines) supporting the redistribution of E-cadherin shown in Figure 6E. In addition, the intensity profiles of the endogenous ZO-1 also showed an increase for iCRT3 treated cells (Supp. Figure 4C, left) which correlated with the increase in ZO-1 staining and Western Blot levels. To assess the functional relevance of the β-catenin signaling pathway in SGEF KD cells, we analyzed the ability of CTRL, SGEF KD, SGEF/Slug dKD, and iCRT3 treated cells to form a monolayer and establish an impermeable barrier, by measuring barrier function using T.E.E.R. Here, we found that inhibiting β-catenin signaling pathway in SGEF KD cells is sufficient to restore the T.E.E.R. to levels comparable to CTRL cells (Figure 6F). On the other hand, T.E.E.R. was not rescued in SGEF/Slug dKD cells, most likely because silencing Slug in SGEF KD cells did not rescue ZO-1 levels (Figure 5A, D). Supporting these findings, we saw a significant increase in the tortuosity of the iCRT3 treated cells, but not in SGEF/Slug dKD cells when compared to SGEF KD cells (Supp. Figure 4D). However, this increased tortuosity in iCRT3 treated cells was still not a complete rescue when compared to CTRL cells tortuosity. These results suggest that the decrease in barrier function observed in SGEF KD can be attributed, at least in part, to the loss of ZO-1.

To summarize, we showed that the expression of E-cadherin and ZO-1 is regulated by a β-catenin mediated signaling pathway. Interestingly, E-cadherin is regulated by Slug, whereas ZO-1 is independently regulated by a yet to be characterized different factor. Finally, we concluded that restoring ZO-1 expression is sufficient to rescue MDCK cells barrier function in the absence of SGEF.

### SGEF regulates junctional E-cadherin mobility

As cell-cell adhesions are established, E-cadherin molecules from neighboring cells initially form weak monomeric *trans* interactions. These monomeric E-cadherin *trans*-pairs can then interact in *cis to* form oligomeric arrays or clusters, which help to strengthen and stabilize the interaction as the AJ matures (Troyanovsky et al., 2021). The ability of E-cadherin to cluster and the dynamics of these clusters can also be regulated by multiple factors, including the interaction of the cadherin complex with the actin cytoskeleton, actomyosin contractility, other *cis*-interactions, and additional adhesion proteins (Troyanovsky et al., 2021; Yap et al., 2015). As a results, the mobility of E-cadherin at mature junctions is very limited when compared to nascent/immature junctions (Adams et al., 1998; de Beco et al., 2009; Sako et al., 1998; Yamada et al., 2005). We hypothesized that the Scribble/SGEF/Dlg1 ternary complex played a role in the regulation of AJ stability and that disrupting the complex would result in increased E-cadherin mobility at the membrane.

To study the diffusion of E-cadherin in CTRL and SGEF KD we tagged endogenous E-cadherin with mScarlet using a CRISPR Knock-In (KI) system (Figure 7A-B) (Bollen et al., 2022). As expected, the expression levels of the endogenous-tagged E-cadherin are significantly lower in SGEF KD cells, which suggests that the KI did not interfere with the normal regulation of E-cadherin expression. We then used fluorescence recovery after photobleaching (FRAP) in CTRL and SGEF KD E-cadherin-mScarlet KI cell lines to study E-cadherin dynamics. Since it is difficult to perform FRAP experiments in cells grown in transwell filters, we adapted a previously described protocol that allowed us to grow cells on the bottom of an inverted filter for 6 days after confluency (Miyazaki et al., 2023). Before the experiment we inverted the filter on a 3D printed support mounted on a glass bottom dish that placed the cells at a distance from the glass that could be imaged with a 63X oil objective. We then photobleached mature junctions and analyzed their recovery. Figure 7C shows representative pre-bleach, bleach, and post-bleach frames from CTRL and SGEF KD cells (see Movies 1 and 2). The fluorescence recovery along junctions was analyzed as a function of time and displayed as kymographs and average recovery graphs (Figure 7D, E). The kymograph analysis showed that in CTRL cells the bleached area remained uniform with no significant fluorescence recovery within the first minute after bleaching, while in SGEF KD cells it recovered rapidly (Figure 7D). FRAP quantification (Figure 7E) confirms that the speed at which the signal recovered after the bleaching event was faster for SGEF KD cells, with a half-time of 106.6 seconds compared to 140.2 seconds for CTRL. In addition, the overall fluorescence recovery showed a striking difference between the immobile fractions of the two cell lines, with CTRL cells having a higher immobile fraction (55.5%) than SGEF KD cells (44.4%). Overall, this data suggests that in SGEF KD cells, the basolateral pool of E-cadherin is more mobile compared to CTRL cells, suggesting E-cadherin is less stable.

**Figure 7.**
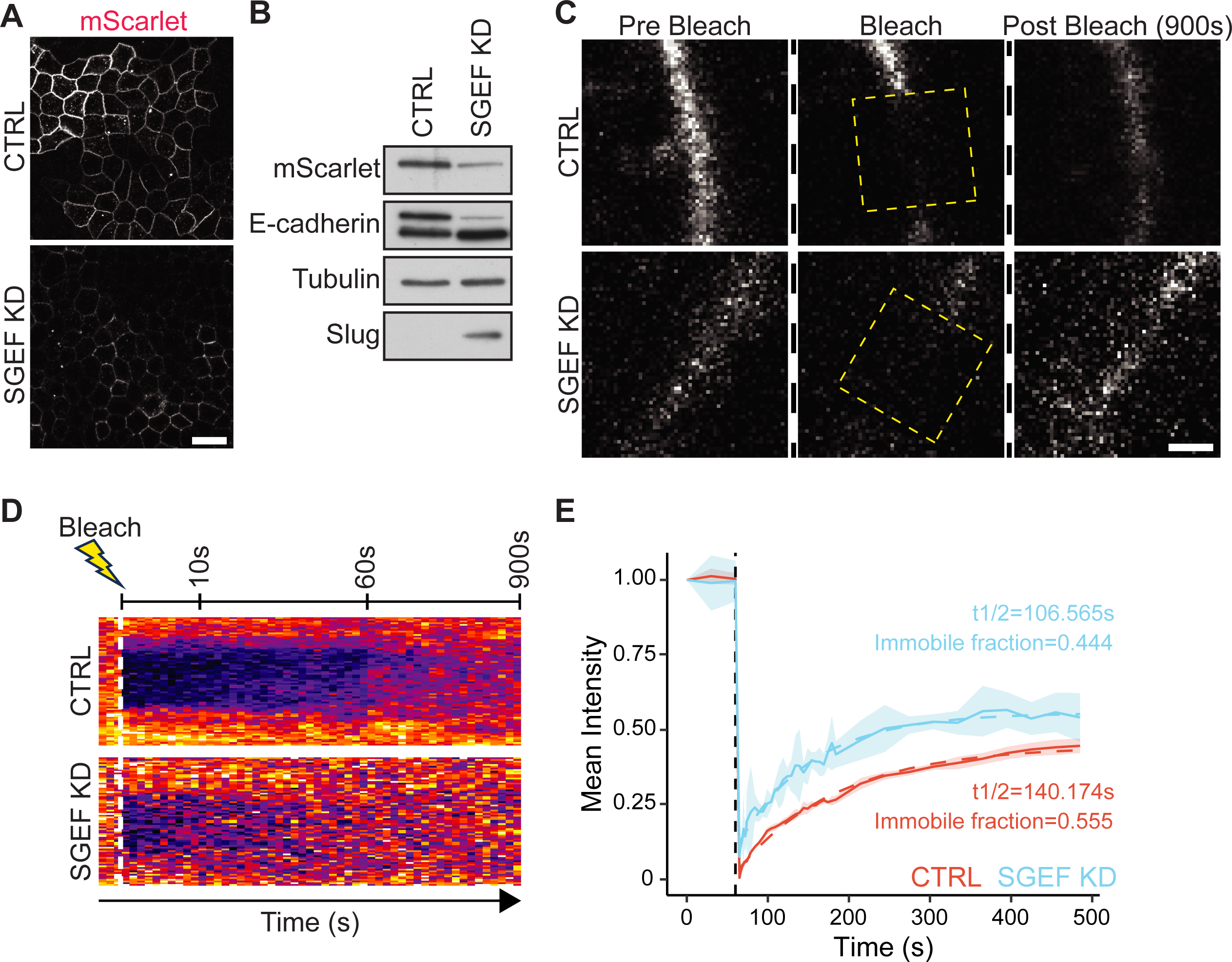
SGEF regulates junctional E-cadherin mobility. **(A)** E-cadherin was endogenously tagged with mScarlet in CTRL and SGEF KD MDCK cells. Representative images for each cell line are shown. Scale bar 20µm. **(B)** Total cell lysates from confluent CTRL and SGEF KD E-cadherin mScarlet KI cells were analyzed by WB for mScarlet and E-cadherin. Tubulin was used as a loading control. **(C)** E-cadherin dynamics was analyzed by FRAP in CTRL and SGEF KD E-cadherin mScarlet KI cells grown for 6 days on inverted filters. Representative frames of a bleached junction for CTRL and SGEF KD cells. The dashed square shows the bleached region. Scale bar 2 µm. **(D)** Kymograph showing the recovery over time for a CTRL and SGEF KD representative junction. **(E)** Fluorescence recovery curves showing the average of multiple junctions across independent experiments (n=4,>35 junctions bleached per condition/experiment). Shaded area represents SD. A regression curve was fitted, and the t_1/2_ of recovery/immobile fraction calculated.

### Silencing SGEF dissociates the E-cadherin/p120-catenin complex and promotes the internalization of E-cadherin

Our results suggest that the members of the Scribble/SGEF/Dlg1 complex may function to stabilize E-cadherin at the membrane, particularly SGEF, which causes the most severe effect when silenced. E-cadherin stability at the basolateral membrane has been shown to be dependent on its interaction with p120-catenin. Dissociation of the E-cadherin/p120-catenin complex or reducing p120 catenin levels results in a significant increase in E-cadherin internalization (Davis et al., 2003; Ireton et al., 2002; Ishiyama et al., 2010).

We did not observe a decrease in p120-catenin or β-catenin levels upon SGEF silencing, but their localization was affected (Figure 1C, E) (Awadia et al., 2019). Based on these results, we hypothesized that silencing SGEF may dissociate the E-cadherin/p120-catenin complex, which would stimulate E-cadherin internalization and the subsequent release of β-catenin, which can then translocate to the nucleus where it regulates transcription. To test this hypothesis, we co-immunoprecipitated the E-cadherin/p120-catenin complex in CTRL and SGEF KD cells, using p120-catenin antibodies and blotting for E-cadherin. Our result showed a significant decrease in the amount of co-immunoprecipitated E-cadherin in SGEF KD cells (Figure 8A). The reciprocal co-immunoprecipitation, E-cadherin immunoprecipitation and blotting for p120-catenin, showed similar results, with a significantly reduced amount of p120-catenin precipitating with E-cadherin in SGEF KD cells (Figure 8B). These results suggest that SGEF plays a role in the formation and/or stabilization of the E-cadherin/p120-catenin complex.

**Figure 8.**
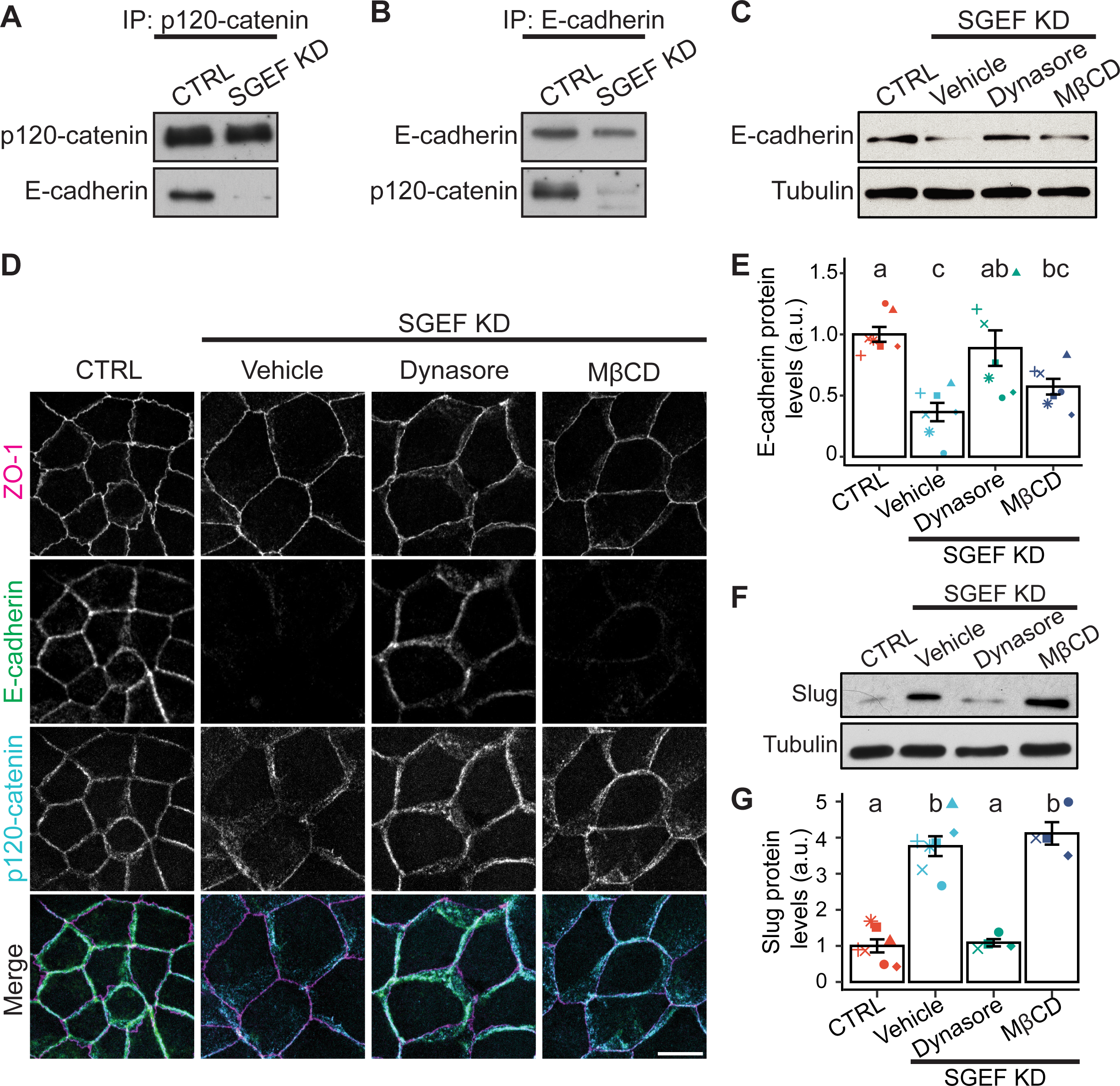
Silencing SGEF dissociates the E-cadherin/p120-catenin complex and promotes the internalization of E-cadherin. **(A-B)** Endogenous p120-catenin (A), or E-cadherin (B), were immunoprecipitated from CTRL and SGEF KD MDCK lysates and immunoblotted for p120-catenin, and E-cadherin. **(C)** Total cell lysates from confluent CTRL (non-treated) and SGEF KD MDCK cells treated with the indicated endocytosis inhibitors for 16 h were analyzed for E-cadherin expression by WB. Tubulin was used as a loading control. **(D)** IF of endogenous ZO-1, E-cadherin and p120 catenin in CTRL (non-treated) and SGEF KD cells after incubation with the indicated endocytosis inhibitors. **(E)** Quantification and ANOVA analysis from WB data in (C) (n=7). Letters indicate Tukey post-hoc significant differences between groups. **(F)** Lysates from CTRL (non-treated) and SGEF KD cells incubated with the different endocytosis inhibitors were analyzed for Slug expression by WB. **(G)** Quantification and ANOVA analysis from WB data in (F) (n=7). The letters indicate Tukey post-hoc significant differences between groups.

One prediction from these results is that E-cadherin internalization should be increased in SGEF KD cells, and that inhibiting endocytosis should restore E-cadherin expression levels. We tested this by treating SGEF KD cells with two different endocytosis inhibitors, dynasore for clathrin mediated endocytosis (CME), and MβCD for clathrin independent endocytosis (Figure 8C, D). Our results showed that dynasore treatment, but not MβCD, restored E-cadherin levels almost completely, confirming the role of endocytosis in the SGEF-dependent downregulation of E-cadherin (Figure 8C-E; Supp. Figure 5C). We also confirmed these results using a different CME inhibitor, pitstop 2 (Supp. Figure 5A).

The results observed after inhibiting CME can be interpreted in two different ways: 1) E-cadherin is still transcriptionally repressed but its levels are rescued after newly synthesized protein accumulates gradually at the membrane when endocytosis is inhibited overnight; or 2) Inhibiting endocytosis prevents the E-cadherin/catenin complex to get internalized, thus preventing β-catenin release, its translocation to the nucleus, and restoring E-cadherin transcription. Using the same inhibitors, we immunoblotted for Slug as a β-catenin signaling indicator. Surprisingly, we saw a striking reduction of Slug expression in the SGEF KD cells treated with dynasore, with levels comparable to CTRL cells, supporting our second hypothesis/scenario (Figure 8F, G). In agreement, immunofluorescence analysis showed that β-catenin is more tightly associated with junctions in dynasore treated SGEF KD cells when compared to the diffuse localization of non-treated SGEF KD cells, or cells treated with MβCD (Supp. Figure 5B, D).

## Discussion

Cell polarity is regulated by the coordinated action of three highly conserved protein complexes; PAR, Crumbs, and Scribble (Bilder et al., 2003). The Scribble complex, which comprises Scribble, Dlg1 and Lgl, was originally identified in Drosophila as a critical regulator of epithelial polarity (Elsum et al., 2012). It was later shown to be involved in the regulation of other cellular processes, including cell-cell adhesion, asymmetric cell division, vesicular trafficking, cell migration, and planar-cell polarity (Elsum et al., 2012). In mammalian cells, the Scribble complex also plays key roles in the regulation of cell adhesion and polarity (Bonello and Peifer, 2019). Importantly, dysregulation of the Scribble complex is commonly observed in human cancers and correlates with tumor progression (Elsum et al., 2012).

Most information to date on the function of the Scribble complex originates from genetic studies in flies, or loss of function experiments in mammals (Elsum et al., 2012). With no known catalytic activity, the proteins in the Scribble complex are believed to function as scaffolding platforms to recruit other binding partners. These include the Rho GTPases and their regulators, such as RhoGEFs and RhoGAPs, which will build spatially distinct signaling complexes (Bonello and Peifer, 2019; Iden and Collard, 2008a; Mack and Georgiou, 2014). However, it is not known which downstream signaling pathways are regulated by the Scribble complex.

Previous work by our lab has shown that SGEF, a RhoG specific GEF, forms a ternary complex with two members of the Scribble polarity complex, Scribble and Dlg1 (Awadia et al., 2019). We also showed that SGEF regulates the assembly and function of AJ and TJ in both 2D and 3D. Notably, silencing SGEF results in a dramatic downregulation of the expression of E-cadherin in epithelial cells. This loss of E-cadherin is relevant because E-cadherin is a central component of cell-cell adhesion and is required for epithelial development in the embryo, and to maintain epithelial homeostasis in an adult (Hartsock and Nelson, 2008). Moreover, the downregulation of E-cadherin expression is an important hallmark of EMT, which can be observed during development and cancer. In addition, E-cadherin is a tumor suppressor gene that, when downregulated, increases invasion *in vitro* and promotes the development of malignant epithelial cancers *in vivo* (Bruner and Derksen, 2018; Thiery, 2002; Thiery and Sleeman, 2006). However, the mechanisms by which this novel Scribble/SGEF/Dlg1 complex regulates E-cadherin stability and expression levels are not known.

In this study, we were able characterize the role of the Scribble/SGEF/Dlg1 ternary complex in the SGEF-mediated regulation of E-cadherin expression and junction integrity. Our work shows that SGEF is the only member of the ternary complex that regulates the expression levels of E-cadherin. However, Scribble and Dlg1 KO cells showed an altered AJ architecture but no significant changes in E-cadherin expression. This suggests that Scribble and Dlg1 are involved in the regulation of E-cadherin stability at the AJ, as it has been suggested in previous literature (Firestein and Rongo, 2001; Ivanov et al., 2010; Lohia et al., 2012; Qin et al., 2005; Yates et al., 2013). Importantly, SGEF is the only member of the complex that has a catalytic function, and we were able to show that the catalytic activity is essential for the regulation of E-cadherin expression (Awadia et al., 2019). Here, we extended these studies to demonstrate that one of the roles of the Scribble/SGEF/Dlg1 complex is to target SGEF activity to the basolateral membrane. Our results showed that expressing the catalytic domain of SGEF, in SGEF KD cells, rescues the expression levels of E-cadherin, but only when it is targeted to the basolateral membrane. This, together with our previous findings, suggests that the activity of RhoG, localized to the basolateral membrane by Scribble and Dlg1, is involved in the regulation of E-cadherin expression (Awadia et al., 2019). Based on these results and previous studies showing that SGEF may exist in an autoinhibited state, we believe its association with the complex may also function to release this autoinhibitory state to activate SGEF on site (Zhang et al., 2021).

Furthermore, we show that all three members of the Scribble/SGEF/Dlg1 ternary complex play a role in the regulation of TJ architecture and barrier function. Knocking out Scribble or Dlg1 phenocopies the changes previously observed in SGEF KD cells, which include larger cell area, decreased aspect ratio, decreased TJ tortuosity, and impaired barrier function (Awadia et al., 2019). In contrast to the downregulation of E-cadherin, the effects on TJ are independent of the catalytic activity of SGEF and appear to be mediated by ZO-1, as silencing either SGEF, Scribble or Dlg1 resulted in a significant decrease of junctional ZO-1. This seems to involve ZO-1 downregulation in SGEF KD cells, and its redistribution in Scribble and Dlg1 KO cells, although more work needs to be done to define the mechanisms. Previous literature has shown that the ZO family of TJ proteins is involved in defining the characteristic “wavy” pattern of MDCK cells and that knocking out members of this family resulted in a straightening of the junctions as well as an increase in permeability (Fanning et al., 2012; Tokuda et al., 2014; Van Itallie et al., 2009). Overall, these results confirm our previous work, where we defined a scaffolding and a catalytic role for the Scribble/SGEF/Dlg1 ternary complex, with the scaffolding function involved in the regulation of TJ permeability and the catalytic role regulating E-cadherin expression. Silencing SGEF impacts both the scaffolding and catalytic roles, and thus results in a more severe phenotype, whereas silencing Dlg1 or Scribble affects mainly the scaffolding role but not the catalytic role (or at least not in a significant manner) (Awadia et al., 2019).

How does SGEF regulate E-cadherin stability and expression? Our results demonstrate that the main driver of E-cadherin downregulation in SGEF KD cells is a decrease in the mRNA levels. We also found that the downregulation of E-cadherin transcription is mediated by the transcriptional repressor Slug, which is significantly upregulated in SGEF KD cells, in a process that is mediated by an increase in the β-catenin signaling pathway. The catalytic activity of SGEF is essential for the regulation of Slug levels, as the catalytic dead mutant cannot rescue the phenotype. In contrast, Scribble and Dlg1 KO cells, which show no significant decrease in E-cadherin levels, displayed only a minor increase in Slug levels in comparison to SGEF KD cells, confirming the key role of the catalytic activity of SGEF in the regulation of E-cadherin levels. Interestingly, silencing Slug in SGEF KD rescues E-cadherin levels, but it is not sufficient to restore its proper localization, supporting our hypothesis that an intact Scribble/SGEF/Dlg1 complex stabilizes E-cadherin at the membrane. Our results also show that proteasomal degradation E-cadherin can potentially contribute to its downregulation, but apparently not at a significant level. With stable SGEF KD we cannot distinguish whether E-cadherin proteasomal degradation occurs early, when SGEF is being actively downregulated, and then stops when transcription is repressed, or if it is not a significant factor. Using inducible SGEF KD to analyze these early time points would help us to answer this question.

The amount of E-cadherin at the plasma membrane is determined by a dynamic equilibrium between synthesis, degradation, endocytosis, and recycling (Kowalczyk and Nanes, 2012). E-cadherin can be endocytosed through different mechanisms, including clathrin-dependent, clathrin-independent, and caveolae-mediated (Bryant and Stow, 2004; Ivanov et al., 2004; Le et al., 1999; Lohia et al., 2012; Qin et al., 2005). Once internalized, E-cadherin can be recycled back to the plasma membrane, sequestered in internal compartments, or sent for degradation (Bryant and Stow, 2004). Endocytosis of E-cadherin is triggered by the disassembly of the E-cadherin/p120-catenin complex (Davis et al., 2003; Ireton et al., 2002; Ishiyama et al., 2010). Our results suggest that the downregulation of E-cadherin levels in SGEF KD cells is initiated by the dissociation of p120-catenin from the E-cadherin complex, followed by internalization of E-cadherin through endocytosis and degradation by the proteasome. Interestingly, inhibiting CME rescued E-cadherin and Slug expression to normal levels, placing the complex dissociation and internalization of E-cadherin as the earliest of a series of events that mediates its downregulation.

One piece of the puzzle that remained unanswered in this study is the identity of the transcription factor controlling ZO-1 expression. Our results show that ZO-1, like E-cadherin, is regulated at the transcriptional level in SGEF KD cells in a process mediated by β-catenin, but in a Slug independent fashion. A potential candidate is ZEB1, a transcriptional repressor also regulated by the β-catenin signaling pathway, that is mostly associated with E-cadherin repression (Eger et al., 2005; Sanchez-Tillo et al., 2011). ZEB1 can also regulate the expression of ZO-1 indirectly via a MAPK-ERK pathway (Liu et al., 2018). Interestingly, Scribble has also been shown to play a role in the regulation of ZEB1 levels (Elsum et al., 2013). However, we have not detected changes in ZEB-1 expression upon SGEF KD, suggesting that in this case, ZO-1 regulation by the Scribble/SGEF/Dlg1 complex is mediated by a different pathway.

In this study we have shown that the activation of RhoG by SGEF at the basolateral membrane is an essential component for the regulation of E-cadherin stability and expression. However, we did not explore the downstream effectors of RhoG. Perhaps the most likely candidate is ELMO2 (engulfment and cell motility 2), which is known to be a specific effector for RhoG (Katoh and Negishi, 2003). ELMO2 has been involved in the recruitment of E-cadherin to initial cell-cell adhesion sites in MDCK cells (Toret et al., 2014a). The SGEF-RhoG axis may be involved in the recruitment of ELMO2, which can function to locally activate Rac1 at the cell-cell adhesion initiaition sites by forming a complex with the Rac1-GEF DOCK1 (dedicator of cytokinesis 1) (Toret et al., 2014a). Interestingly, ELMO2 has also been shown to function in Rab11-mediated recycling of E-cadherin containing endosomes in keratinocytes, together with the Integrin Linked Kinase (ILK) (Chen et al., 2013; Tan et al., 2001; Wu et al., 1998). In addition, ELMO2, RhoG and ILK can form a ternary complex (Ho and Dagnino, 2012; Jackson et al., 2015). Finally, silencing ELMO2 affects E-cadherin localization and expression levels, but not as drastically as silencing SGEF (Toret et al., 2014a; Toret et al., 2014b). This may be a result of residual ELMO2 expression due to incomplete silencing, or alternatively suggest that SGEF and RhoG modulate more than one downstream output, which could include ELMO2 as well as other yet to be characterized effectors. Our previous work has shown that silencing RhoG increases Src activation (Goicoechea et al., 2017). Interestingly, Src-dependent phosphorylation of the E-cadherin at the juxta membrane domain disrupts E-cadherin binding and pormotes its internalization (Fujita et al., 2002). Future efforts will be devoted to dissecting the pathway that controls E-cadherin dynamics and turnover downstream of SGEF and RhoG.

An interesting observation from this study was that rescuing E-cadherin without restoring SGEF’s catalytic activity, e.g., SGEF/Slug dKD or iCRT3 treatment, was not sufficient to restore E-cadherin proper localization at AJs and suggests that SGEF plays a role stabilizing E-cadherin at the membrane.

As intercellular junctions mature, E-cadherin/catenin complexes in AJs form clusters driven by *cis-* and *trans-* interactions in the cadherin ectodomain. The stability of these extracellular clusters is further enhanced by the binding of α-catenin to actin filaments and through interaction with proteins associated with the E-cadherin/catenin complex (Mege and Ishiyama, 2017; Troyanovsky, 2023; Yap et al., 2015). As E-cadherin assembles into these clusters it becomes less mobile at the membrane, showing little to no membrane diffusion along mature junctions, and most of E-cadherin’s turnover in stable junctions can be attributed to endocytosis and recycling (de Beco et al., 2009). However, FRAP experiments have demonstrated than inhibiting *cis-* and/or *trans-* interactions increases E-cadherin mobility at the membrane, reduces the immobile fraction and destabilizes junctions (Erami et al., 2015; Harrison et al., 2011; Strale et al., 2015).

Our results show that E-cadherin localization at junctions is more diffuse in cells where the Scribble/SGEF/Dlg1 complex is disrupted, which suggests it is less stable and potentially more mobile at the membrane. This was confirmed in FRAP experiments where we followed the dynamics of endogenously tagged E-cadherin in CTRL vs SGEF KD cells. Our data shows that E-cadherin has a faster diffusion/exchange and lower immobile fraction in SGEF KD cells and provides support to our hypothesis. Recently, Troyanovsky and colleagues showed that Scribble can mediate the formation of a specific population of E-cadherin clusters, which are different from the E-cadherin/catenin clusters formed at AJs and stabilized by the interaction between α-catenin and actin (Troyanovsky et al., 2021). This is supported by previous work showing faster FRAP recovery of E-cadherin-GFP when Scribble is silenced (Lohia et al., 2012). The formation of this Scribble containing clusters is proposed to involve *cis*-interactions between E-cadherin complexes, which are mediated by complaex associated proteins like Scribble (Troyanovsky et al., 2021). For example, two of the four PDZ domains of Scribble have been shown to interact with catenin (Ivarsson et al., 2014; Zhang et al., 2006).

Alltogether, these findings align with our results, supporting the role of the Scribble/SGEF/Dlg1 complex in AJ stability and TJ function. In addition we describe in detail the mechanism by which SGEF’s activity regulates E-cadherin expression levels (Figure 9). Future work will focus in uidnerstanding the mechanisms that control SGEF activation at junctions, the spatiotemporal regulation of RhoG activity during junction formation and the signaling pathwaty downtream of SGEF/RhoG activation.

**Figure 9.**
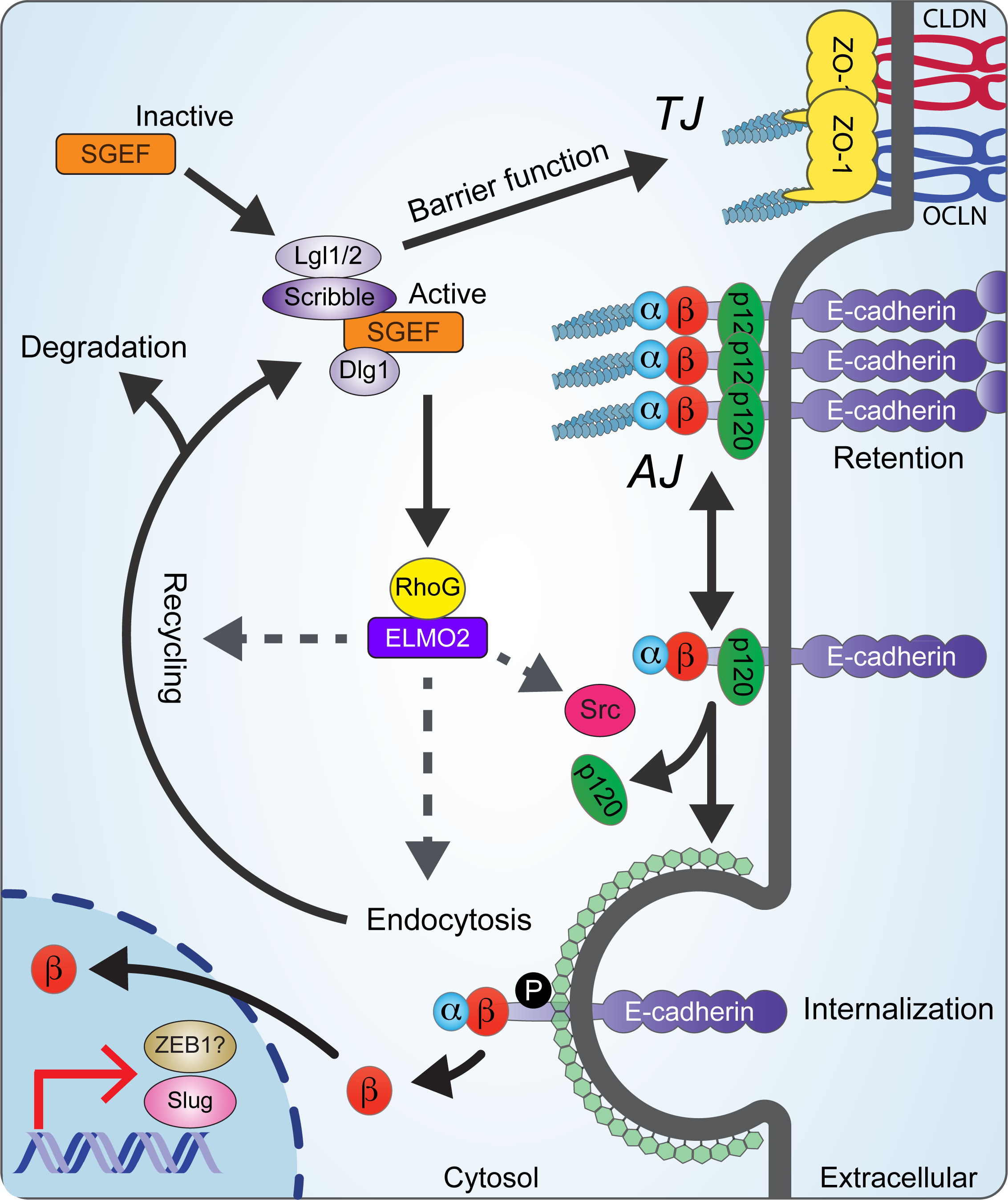
Working model. The Scribble/SGEF/Dlg1 ternary complex controls E-cadherin stability at the membrane. This process requires the catalytic activity of SGEF, which needs to be recruited to the basolateral membrane by Scribble/Dlg1. Downstream of SGEF, RhoG can potentially play a role in different processes, including the phosphorylation of E-cadherin/catenin complex by Src, and E-cadherin recycling and recruitment to membranes through its effector ELMO2. When the complex is perturbed, particularly the activity of SGEF, the E-cadherin/p120-catenin complex dissociates. promoting E-cadherin endocytosis and its eventual degradation. This process releases β-catenin, which can then translocate to the nucleus and induces the transcriptional repressor Slug and other factors, resulting in the downregulation of E-cadherin 1ranscription.

## Materials and methods

### Cell lines

MDCK II cells were a gift from I.G. Macara (Vanderbilt University, Nashville, TN). MDCK cells were grown in DMEM (GIBCO) containing 10% FBS and antibiotics (penicillin-streptomycin). Cell lines were grown at 37°C and 5% CO2. All experiments were conducted with early passage cells that were passaged no more than 20 times. Mycoplasma was tested regularly by staining with Hoechst 33342 (AnaSpec Inc).

### Antibodies and Reagents

The following commercial antibodies were also used throughout this study: RhoGDI (SC-360, rabbit polyclonal) 1:10000 WB from Santa Cruz Biotechnology; tubulin (T9028, mouse monoclonal) 1:50,000 WB, from Sigma-Aldrich; E-cadherin (24E10, rabbit mAb) 1:1,000 WB, 1:200 IF, Slug (C19G7 rabbit mAb) 1:500 WB, Snail (C15D3 rabbit mAb) 1:500 WB, ZEB1 (D80D3 rabbit mAb) 1:500, all from Cell Signaling Technology; ZO-1 (339100, mouse monoclonal) 1:1,000 WB, 1:100 IF, Alexa Fluor 488– and Alexa Fluor 594-conjugated anti-mouse-IgG and anti-rabbit-IgG secondary antibodies (A11008, A11001, A11005, A32733, and R37117), and Alexa Fluor 647 (A22287) conjugated to phalloidin, from Thermo Fisher Scientific; HRP-conjugated anti-mouse-IgG, and anti-rabbit-IgG secondary antibodies from Jackson Immunoresearch (715-035-151, 711-035-152).

The following Reagents were also used throughout this study. Incubation was done ON (∼16h) unless indicated otherwise. HGF was acquired from R&D (P14210) and used at 100 ng/ml. For endocytosis experiments dynasore Hydrate (Sigma-D7693), Pitstop 2 (MedChemExpress-HY-115604/CS-0103973), and MβCD (Sigma-C4555) were used at 100µM, 25 µM, and 2 mM, respectively. For degradation experiments (R)-MG132 (Cayman Chemical-13697) and Chloroquine diphosphate salt (Sigma-Aldrich) were used at 20 µM and 25 µM, respectively. For β-catenin inhibition iCRT3 (EMD Millipore-219332) was used at 50 µM.

### Constructs and Primers

CTRL and SGEF KD MDCK cell lines were established in our previous publication (Awadia et al., 2019). Constructs to KD Slug in SGEF KD were cloned into pLKO.1-Hygro (Addgene 24150). To generate Scribble and Dlg1 CRISPR/Cas9 KO cell lines we used pLentiCRISPR v2-Blast (Addgene 83480). DH-PH and VSV-G-DH-PH Rescue constructs were cloned using gateway recombination (Thermo Fisher) into pLenti CMV Hygro DEST (Addgene 17454). We generated stable cells lines using these constructs via lentiviral delivery and antibiotic selection. For KO cell lines single cell clones were identified and validated. Details for the constructs used in this study, including shRNA and gRNA targeting sequences utilized are listed in Supp. Table 1

### Lentiviral particles and infection

Lentiviral particles were packed using HEK293FT and a standard calcium phosphate transfection method as described previously in (Awadia et al., 2019). Packaging plasmids pMD2.G, pSPAX2, and the lentivirus plasmid were mixed in a molar ratio 1:1:1 (total 10 µg). pMD2.G and pSPAX2 were a gift from Didier Trono, EPFL, Lausanne, Switzerland (Addgene plasmids 12259 and 12260). HEK293FT culture medium was changed 24 h after transfection, and lentivirus particles were harvested 48 h after transfection.

MDCK cells were infected with lentivirus particles overnight. The following day, the infection medium was replaced with complete medium and 500 ng/ml hygromycin to select transformed cells. For some viruses, single-cell colonies were isolated by serial dilution and tested for the corresponding phenotype.

### E-cadherin KI

To tag the endogenous E-cadherin gene (*cdh1*) from MDCK cells we followed the protocol described by Bollen and collaborators (Bollen et al., 2022). The gRNA used was “GAGGTGGCGAGGACGACTAG” which targets the region between D882 and the STOP Codon at the C-terminal portion of E-cadherin. This gRNA was cloned into a Cas9 D10A backbone (PX462) (Addgene 62987). The upstream and downstream homology arms, both 500 bp, were cloned into the plasmid TVBB C-term-mScarlet (Addgene 169219) with the purpose of knocking in mScarlet in frame with D882. Both constructs were then electroporated into MDCK cells using the Neon^TM^ transfection system (Thermo Fisher) and single cell clones isolated by serial dilution and checked for the fluorescent phenotype.

### Real Time PCR

Cells cultured for 4 days at a starting density of 7 x 10^4^/cm^2^ were lysed in TRIzol (Ambion) for 5 min and transferred to an RNA-free microcentrifuge tube following manufacturer recommendations. Cell lysates were clarified, and then total RNA extracted with the Direct-zol RNA miniPrep kit (Zymo). cDNA was synthesized using Superscript IV RT polymerase (Invitrogen), RNA pol inhibitor (Applied Biosystems), and 2000 ng of RNA as indicated by the manufacturer. Finally, real time PCR was carried out using 100-200 ng of template, 200 nM of primers and Power SYBR™ Green PCR Master Mix (Applied Biosystems) in a BioRad thermocycler. Ct’s were then normalized by ΔΔCT formula and relative expression compared across at least 3 independent experiments. For this analysis we used student t-test as ΔΔCT normalization lacks variability for CTRL conditions and an ANOVA analysis could be misleading.

### PAGE and Western Blotting

Cells cultured for 4 days at a starting density of 7×10^4^/cm^2^ were rinsed with PBS and then scraped into a lysis buffer containing 50 mM Tris-HCl, pH 7.4, 10 mM MgCl2, 150 mM NaCl, 1% Triton X100, and EZBlock protease inhibitor cocktail (BioVision). The supernatant was collected after centrifugation at 20,000 *g* for 10 min. For immunoblotting, lysates were boiled in 2X Laemmli buffer, and 20-60 μg of protein was resolved by SDS-PAGE. The proteins were transferred onto PVDF membranes and immunoblotted with the indicated antibodies. Immunocomplexes were visualized using the SuperSignal West Pico PLUS Chemiluminescent HRP substrate (Thermo Fisher Scientific).

### T.E.E.R. measurements

To measure T.E.E.R., the cells were plated onto a 0.4 μm polyester membrane, at 1 × 10^5^/cm^2^ (Corning). Resistance measurements were performed 4 days later in triplicate using an EVOM Epithelial Volt/ohm meter (World Precision Instruments). The values represent the average of three replicates minus the background resistance. Values were normalized to those of CTRL cells (100%).

### Immunofluorescence assays and Microscopy

The immunofluorescence (IF) assays were performed as described before (Garcia-Mata et al., 2003). Briefly, MDCK cells grown on coverslips or Transwell filters (Corning) were fixed for 10 min with 4% paraformaldehyde and quenched with 10 mM ammonium chloride. Cells were then permeabilized with 0.1% Triton X-100 in PBS for 10 min. The coverslips were then washed with PBS and blocked with PBS, 2.5% goat serum, 0.2% Tween 20 for 5 min, followed by 5 min of blocking with PBS, 0.4% fish skin gelatin, 0.2% Tween 20. Cells then were incubated with the primary antibody for 1 h at room temperature. Coverslips were washed five times with PBS, 0.2% Tween 20 followed by 5 min of blocking with PBS, 0.4% fish skin gelatin, 0.2% Tween 20 and 5 min with PBS, 2.5% goat serum, 0.2% Tween 20. Secondary antibody diluted in the blocking solution was then added for 45 min, washed five times with PBS, 0.2% Tween, and mounted on glass slides using ProLong Diamond Antifade Mountant (P36965; Thermo Fisher Scientific).

Confocal images were collected using a Leica Stellaris 5 laser scanning confocal microscope equipped with an HC PL APO 63X/1.40 OIL CS2 objective, HyD detectors and a Tokai Hit STX stage top incubator (set to 37°C and 5% CO2 for live imaging).

### Image processing, segmentation, and quantification

All quantification of junctional E-cadherin and ZO-1 was carried out on confocal Z-stacks acquired under Nyquist parameters with a HC PL APO 63X/1.40 OIL CS2 objective. Two different approaches were used in this study: a 3D depth analysis of junctional proteins (as Figure 5F), and a perpendicular analysis of junctional intensity (as Figure 2E).

For segmentation of MDCK monolayers we used the TJ marker ZO-1. Briefly, we used the plugin Tissue Analyzer from ImageJ (Aigouy et al., 2010) and a 3 µm Max projection aligned using the brightest point of ZO-1 staining as reference. Segmentation was manually corrected, and junctions exported as regions of interests (ROIs) to the ImageJ ROI manager.

For the 3D depth analysis junctions segmented with Tissue analyzer as stated before were exported to the ImageJ ROI manager. The following steps were carried out as part of MACRO script in ImageJ. Individual ROI was widened (∼40 px) and transformed to Area (LineToArea) and this was used to measure each Z-plane for each staining. Measurements were saved by staining and junction into different CSVs. From here onwards data was wrangled using RStudio and the Tidyverse package. Raw data from all the ROIs measured were imported to RStudio and then aligned to the brightest Z-plane for ZO-1. Data was normalized for each experiment using the min-max feature scaling and graphed using the ggplot library. At least 3 fields from at least 3 independent experiments were used for quantification unless otherwise indicated.

For the perpendicular analysis of junctional intensity, we used ImageJ and did a 3µm apical projection defined by the brightest point of Actin staining. Then, we manually drew perpendicular lines to the junctions (40px wide) and measured the intensity for Actin, E-cadherin, or ZO-1. As before, we used RStudio and the Tiddyverse package to wrangle the data. The brightest point of actin was used to center all measurements and avoid bias of the other proteins. Data was normalized for each experiment using the min-max feature scaling and graphed using the ggplot library. At least 3 fields from at least 3 independent experiments were used for quantification unless otherwise indicated.

For the tortuosity index measurements, we defined tortuosity index as the coefficient between the cell perimeter and the minimum polygon connecting the cell tricellular junctions. To achieve this, we segmented cells using the plugin Tissue Analyzer as described above and exported whole cell segmentations as ImageJ ROIs. Then, we manually traced the minimum polygon connecting tricellular junctions for each cell.

### Fluorescence recovery after photobleaching (FRAP)

To measure E-cadherin dynamics in live cells, we cultured cells in inverted filters following the guidelines from Miyazaki and collaborators (Miyazaki et al., 2023). MDCK cells expressing E-cadherin::mScarlet were plated onto inverted 0.4 μm polyester membrane, at 1 × 10^5^/cm^2^ (Corning), and they were incubated for ∼2h until they attached. Then, filters were placed in their corresponding vessel and grown for 6 days with media being changed every other day. To be able to image the inverted filters we designed a spacer which was 3D printed and could be sterilized and reused. This spacer fits in an Ibidi 35 mm glass bottom µ-dish and allows the cells to be close to the glass bottom for FRAP microscopy. FRAP data was collected using the FRAP module LAS X software. 3 Frames every 30s were collected pre bleach. Post bleach acquisition was done in 3 different timeframes. The first 10 seconds were imaged every second (10 frames). The next 50 seconds were imaged every 5 seconds (10 frames). Lastly, we continued imaging every 30 seconds until the experiment reached 15 minutes (25 frames). For bleaching laser intensity was kept at 90% and multiple bleach events had to be done for proper signal reduction.

FRAP data was then analyzed using RStudio and the Tiddyverse library. At least 35 junctions per condition per experiment, from 4 independent experiments were analyzed. The data was then normalized using the min-max feature scaling, a mathematical regression calculated to get the immobile fraction and t_1/2_ recovery and graphed using the ggplot library.

### Statistical analysis

Technical replicates are defined in this study as individual junctions/cells (IF) or pipetting replicates (qPCR) while biological replicates are defined as the averages of technical replicates from independent experiments. Western blot data does not have technical replicates, only biological replicates. Error bars represent the standard error of the mean (SEM) unless otherwise indicated.

Statistical analysis was done in RStudio using the processed data and either student t-test or ANOVA with a Tuckey post-hoc analysis. Data distribution was assumed to have normal distribution, (Fay and Gerow, 2013) but this was not formally tested. P < 0.05 (*) was considered statistically significant, P < 0.005 (**), and P < 0.001(***).

**Table S1:**
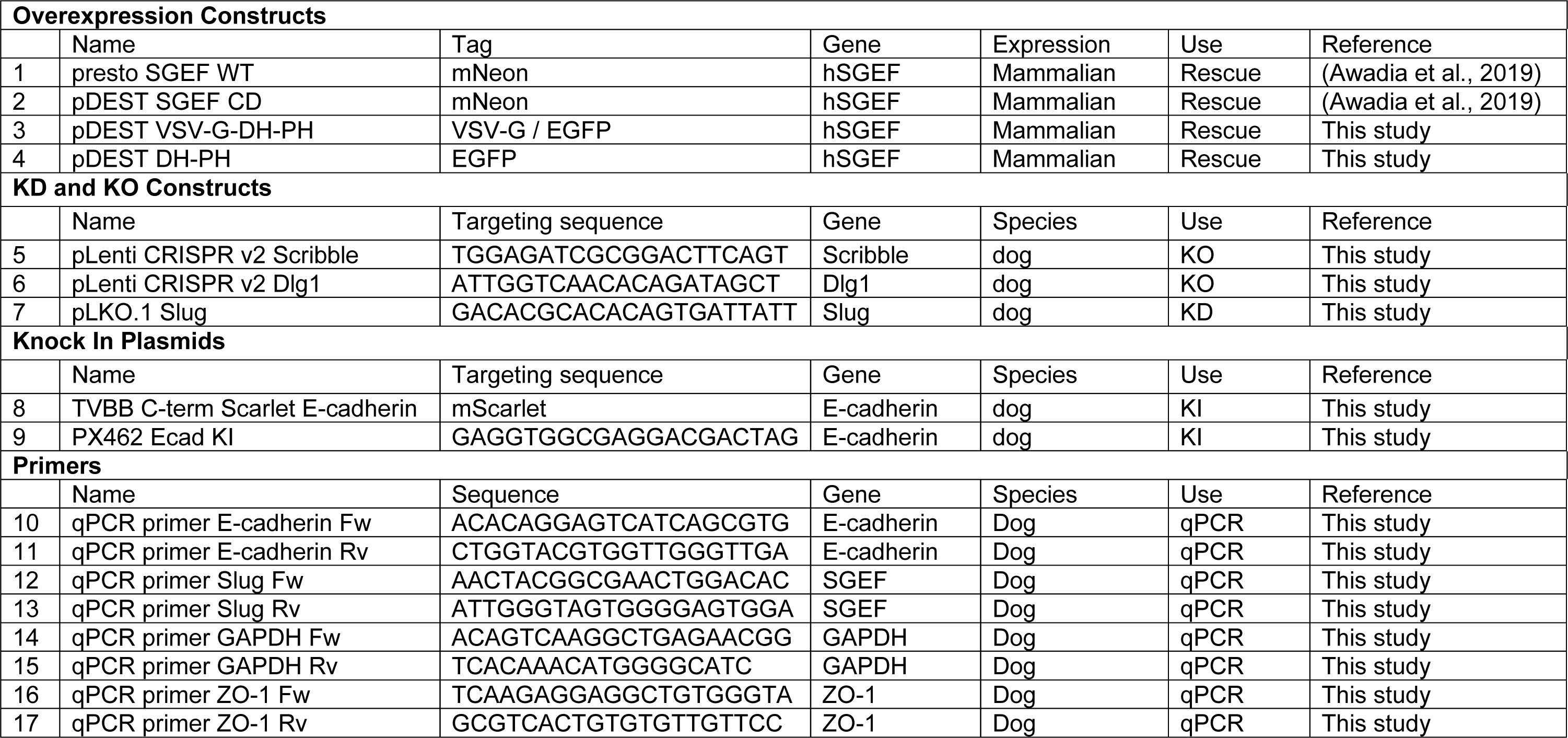
Constructs and primers used in this study.

## Figure Legends

**Supplementary Figure 1.**
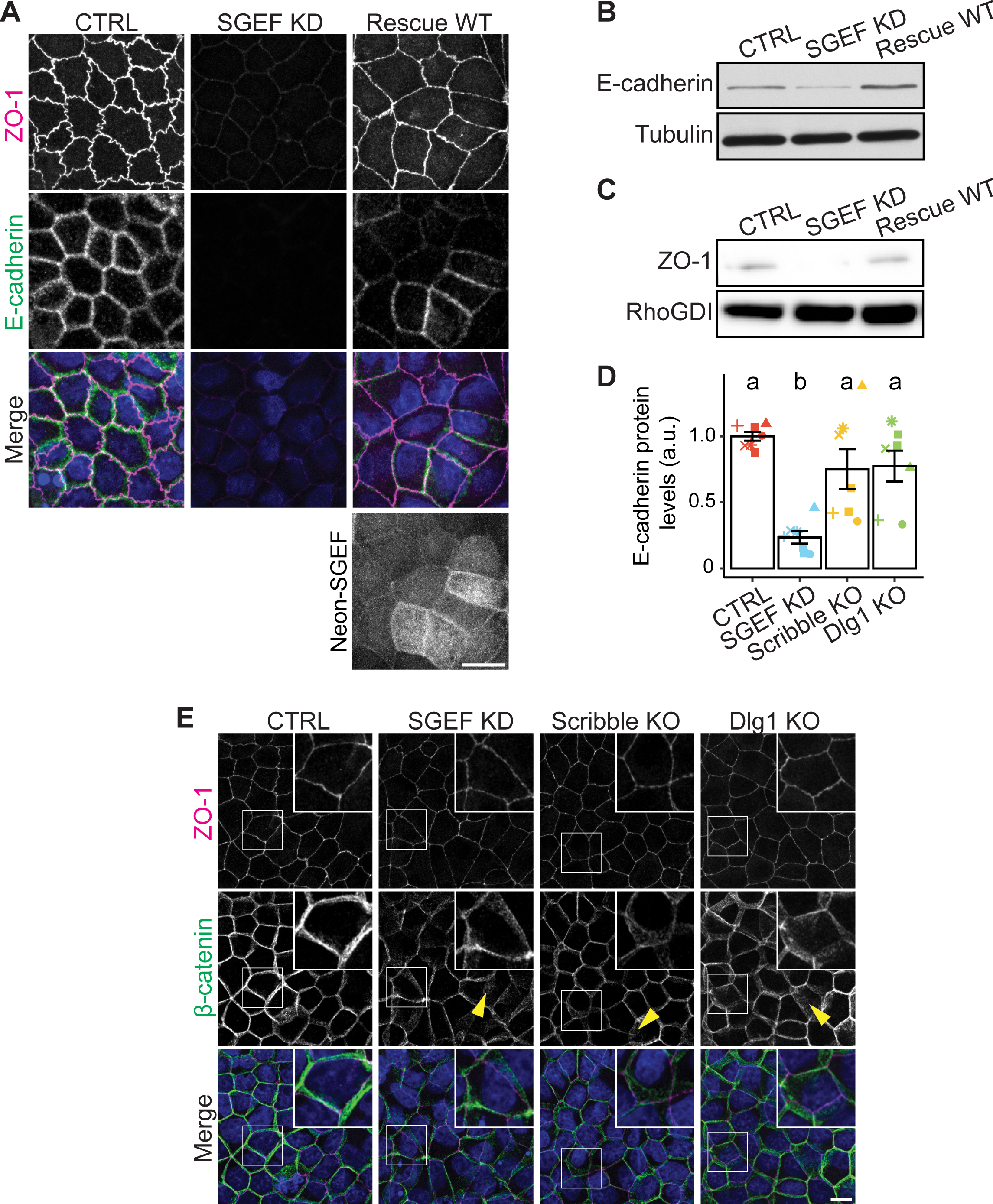
**(A)** IF showing the distribution of ZO-1 and E-cadherin in CTRL, SGEF KD, and SGEF KD cells rescued with SGEF WT grown in permeable filters. All images are 3 µm max projections of the subapical domain (ZO-1 signal used for centering). Scale bar 10 µm. **(B-C)** Total cell lysates from confluent CTRL, SGEF KD, and SGEF KD cells rescued with SGEF WT were analyzed by WB for E-cadherin (B) and ZO-1 (C). Tubulin was used as loading control. **(D)** Quantification and ANOVA analysis from WB results in Figure 1A. The letters indicate Tukey post-hoc significant differences between groups (n=7). **(E)** IF showing the distribution of ZO-1 and β-catenin in CTRL, SGEF KD, and SGEF KD cells rescued with SGEF WT. The inset shows a zoomed image of the region indicated by a white square. All images are 3 µm maximum projections of the subapical domain (ZO-1 signal used for centering). Scale bar 10 µm.

**Supplementary Figure 2.**
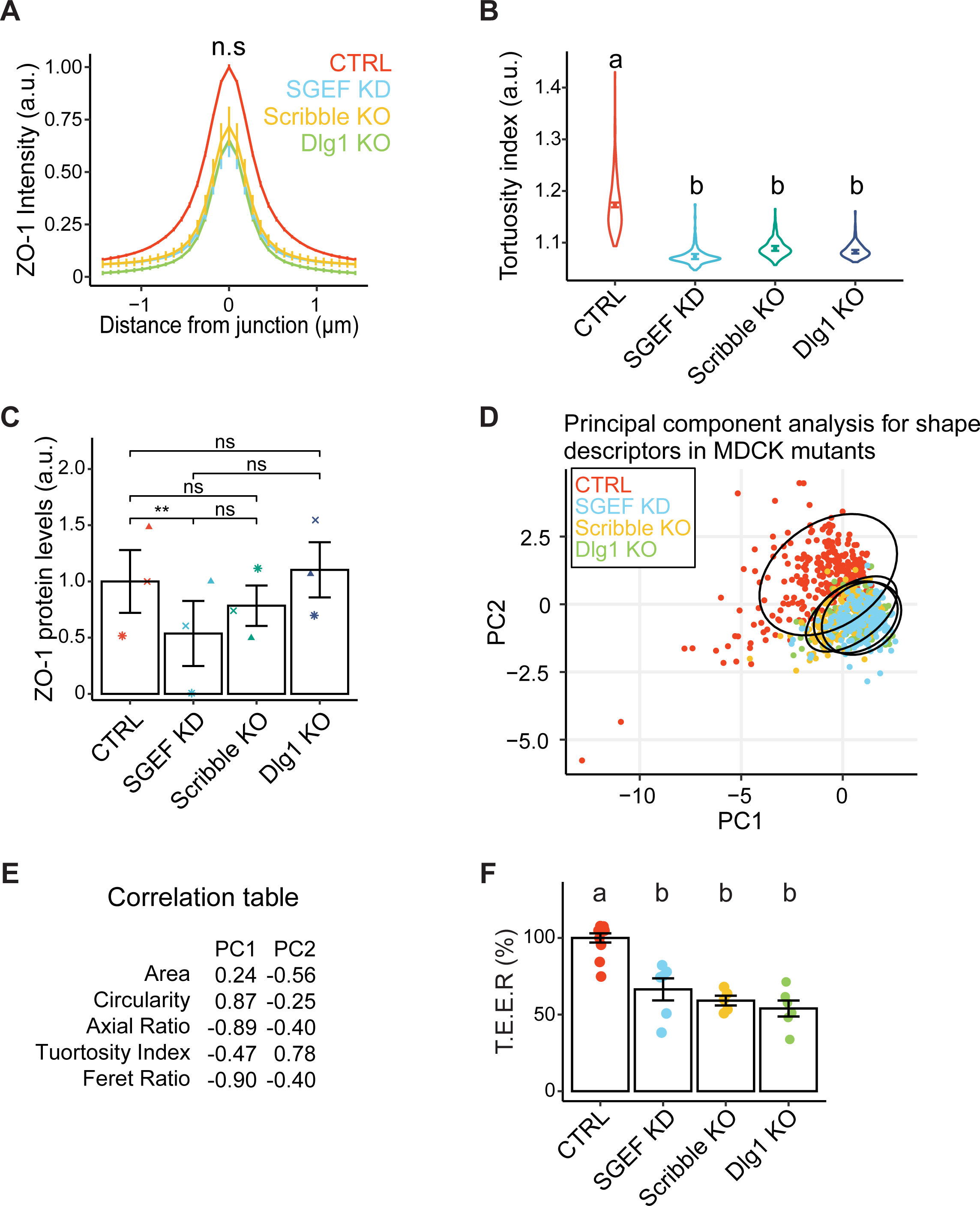
**(A)** Quantification of ZO-1 intensity measured from a perpendicular line across the junctions in Fig 1C (n=3; >150 junctions/condition across all experiments). **(B)** Tortuosity index (n=3; >150 cells/condition across all experiments). **(C)** Quantification and paired t-student analysis from WB results in Figure 1B (n=3). **(D)** Principal component analysis (PCA) for shape descriptors in CTRL, SGEF KD, Scribble KO and Dlg1 KO MDCK cells. Each point in the plot represents the values from a single cell. Ellipses comprise the 95% confidence region. **(E)** Correlation table for shape descriptors and tortuosity index with PC1 and PC2. Table shows that main contributors for PC1 are circularity (positive correlation), and axial ratio and feret ration (negative correlation); for PC2, the main contributor is tortuosity index (positive correlation). **(F)** T.E.E.R. was measured in CTRL, SGEF KD, Scribble KO and Dlg1 KO MDCK cells grown in permeable filters. The graph represents the difference in electrical resistance as a percentage of CTRL cells (n=6).

**Supplementary Figure 3.**
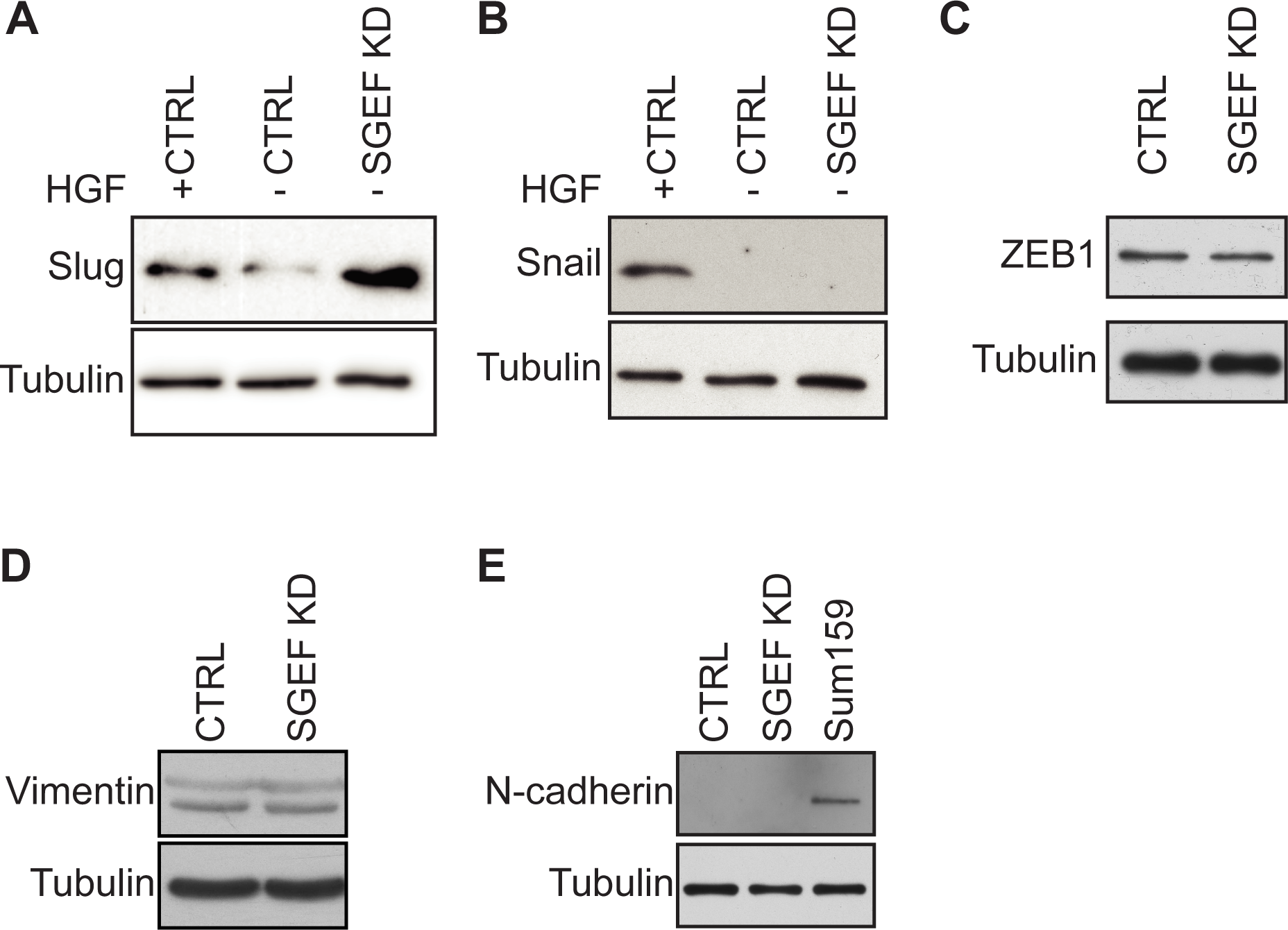
**(A-B)** Total cell lysates from CTRL and SGEF KD MDCK cells were analyzed by WB for the transcriptional repressors Slug (A), Snail (B). HGF was used as a positive control as it promotes EMT in normal epithelial cells. Tubulin was used as a loading control. **(C-E)** Total cell lysates from CTRL and SGEF KD MDCK cells were analyzed by WB for ZEB1 (C), vimentin (D) and N-cadherin (E). Tubulin was used as a loading control. The mesenchymal cell line SUM159 was used as a positive control for N-cadherin.

**Supplementary Figure 4.**
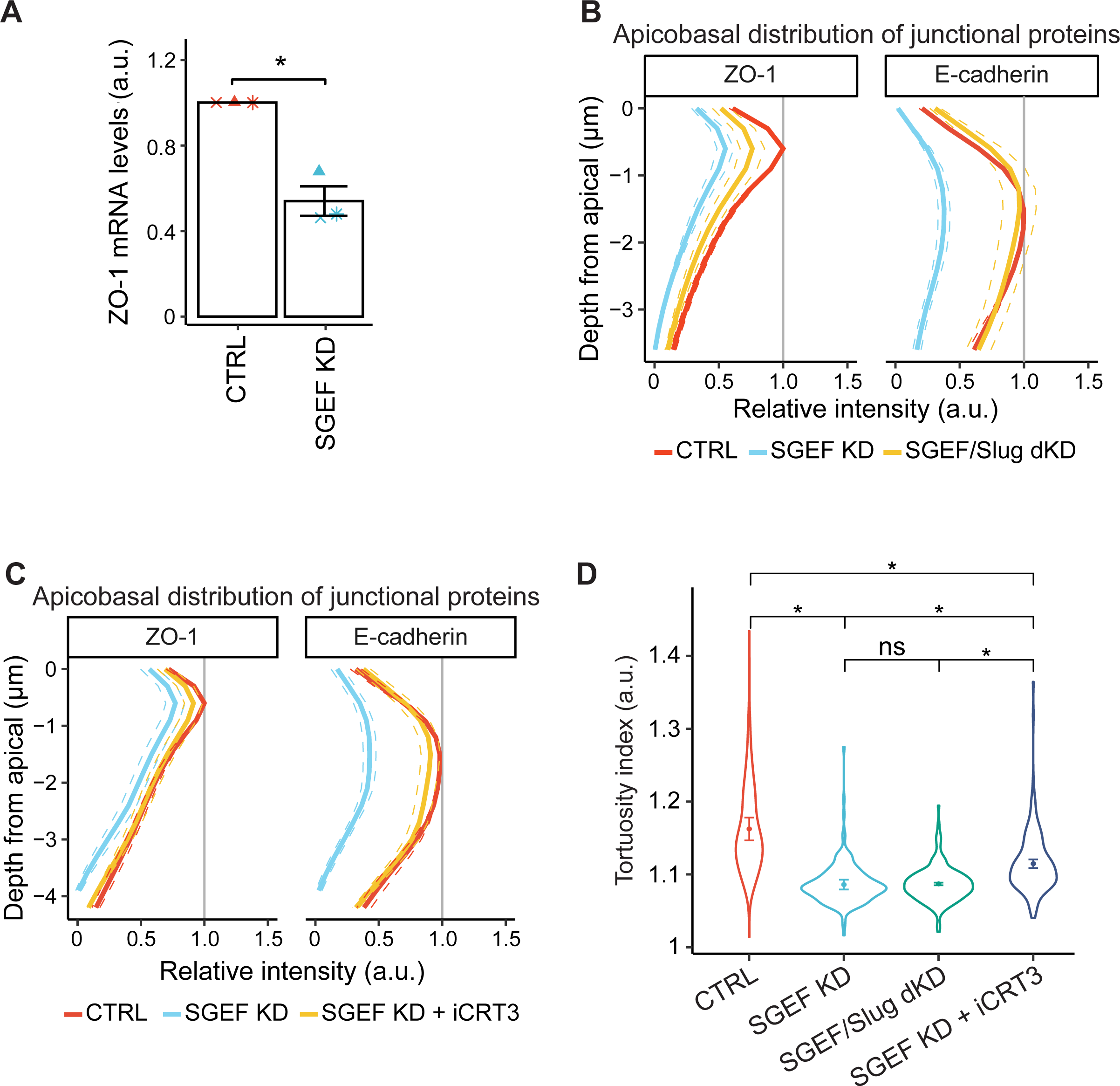
**(A)** qPCR analysis of ZO-1 transcript levels in confluent CTRL, and SGEF KD MDCK cells. **(B)** Quantification of ZO-1 and E-cadherin fluorescence intensity across the depth of the monolayer of CTRL, SGEF KD and SGEF/Slug dKD cells (Figure 5E). (n=5; >1100 junctions/condition across all experiments). Scale bar 10µm (XY) and 5µm (XZ). **(C)** Quantification of ZO-1 and E-cadherin fluorescence intensity across the depth of the monolayer of CTRL, SGEF KD and iCRT3 treated SGEF KD cells (Figure 6E). (n=7; >600 junctions/condition across all experiments). **(B)** Tortuosity index for CTRL, SGEF KD, SGEF/Slug dKD and iCRT3 treated SGEF KD cells. All cells were grown in permeable filters for 6 days (n=4; >150 cells/condition across all experiments).

**Supplementary Figure 5.**
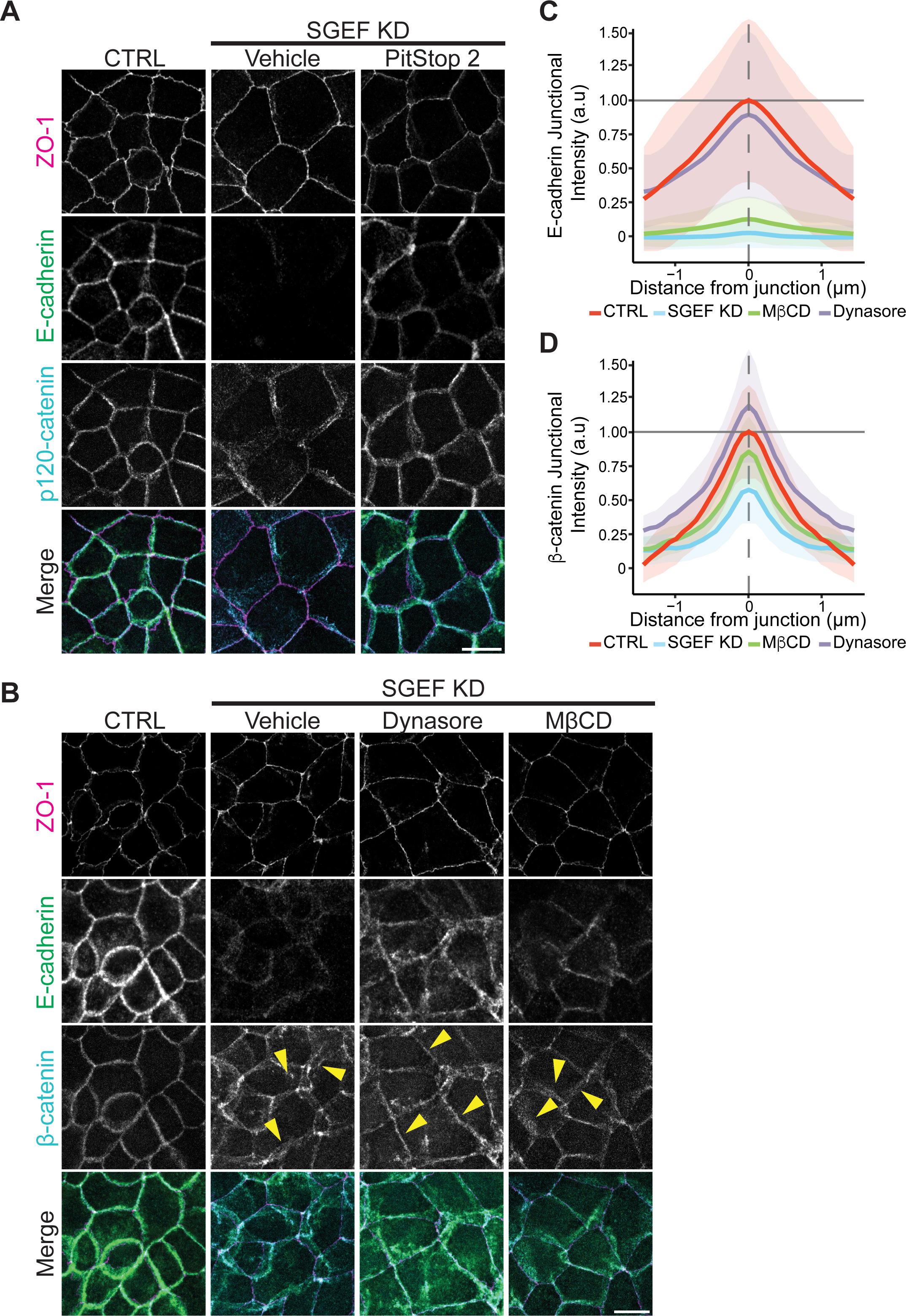
**(A)** IF of endogenous ZO-1, E-cadherin, and p120-catenin in CTRL (non-treated) and SGEF KD cells after incubation with the endocytosis inhibitor PitStop2. Scale bar 10 µm. **(B)** IF of endogenous ZO-1, E-cadherin, and β-catenin in CTRL (non-treated) and SGEF KD cells after incubation with the indicated endocytosis inhibitors. Scale bar 10 µm. **(C)** E-cadherin junctional intensity from (B). (n=1 >30 junctions/condition, shaded area represents SD) **(D)** β-catenin junctional intensity from (B) (n=1; >30 junctions/condition, shaded area represents SD).

**Video 1.** E-cadherin dynamics in live MDCK CTRL cells. CTRL cells expressing endogenous E-cadherin-mScarlet were subjected to FRAP microscopy. Pre-bleach frames were collected every 30 sec. Post-bleach acquisition was done in 3 different timeframes. The first 10 seconds were imaged every second (10 frames). The next 50 seconds were imaged every 5 seconds (10 frames), and the last 15 minutes were imaged every 30 seconds. Arrowheads show bleached regions (frame rate: 24 frames/second).

**Video 2.** E-cadherin dynamics in live MDCK SGEF KD cells. SGEF KD cells expressing endogenous E-cadherin-mScarlet were subjected to FRAP microscopy. Pre-bleach frames were collected every 30 sec. Post-bleach acquisition was done in 3 different timeframes. The first 10 seconds were imaged every second (10 frames). The next 50 seconds were imaged every 5 seconds (10 frames), and the last 15 minutes were imaged every 30 seconds. Arrowheads show bleached regions (frame rate: 24 frames/second).

## Aknowledgements

The authors would like to thank Martijn Gloerich and Kai Simmons for providing reagents. We would also like to thank the members of the Furuse lab and Yildirim-Ayan lab for helping with the FRAP setup. Lastly, we like to thank Andrei Ivanov, Stephan Huveneers, Andrew Goryachev and the members of the Garcia-Mata lab for critical comments and valuable discussions. This work was supported by a grant from the National Institutes of Health to R. Garcia-Mata (R01GM136826).

